# Spiking and locus coeruleus input drive predictive visuomotor plasticity in V1 layer 2/3 neurons

**DOI:** 10.64898/2026.03.03.705098

**Authors:** Scott Yun Ye, Maria Banqueri, Rebecca Jordan

**Affiliations:** Institute for Neuroscience and Cardiovascular Research, University of Edinburgh, 1 George Square, EH8 9JZ, Edinburgh, United Kingdom; Simons Initiative for the Developing Brain

## Abstract

Perception relies on sensory predictions that are continually refined by experience. Prediction errors are neuronal signals thought to instruct plasticity that improves the match between predictions and sensory inputs, minimizing future errors. This central tenet of cortical predictive processing, however, remains untested. Here, we show that activity in two sources of prediction errors – layer 2/3 neurons and locus coeruleus (LC) axons – drives changes in visuomotor integration in mouse V1 consistent with prediction error minimization. Using closed-loop manipulations of layer 2/3 neuron spiking during visuomotor experience, we reveal firing-rate-dependent changes in locomotion-related inputs that increase their opposition with visual inputs. Activation of LC axons facilitated similar activity-dependent reorganization evident in layer 2/3 calcium activity, alongside reduced responses to self-generated visuomotor input. Finally, we demonstrate that visual cortical activity itself can elicit pupil-linked neuromodulation, pointing to a feedback pathway linking cortical output to neuromodulatory plasticity signals. Our results support an activity-dependent, neuromodulation-facilitated cortical mechanism that sharpens visuomotor prediction error computation.

## Introduction

Predicting sensory outcomes, such as those caused by self-motion, reduces redundant information and allows self-generated and externally-generated sensory inputs to be distinguished (Crapse and Sommer, 2008). Because the motor system and the environment are subject to continual change, the learning of predictions must continue throughout life. Predictive processing offers a mechanism for this: sensory predictions are continually updated by prediction errors – neuronal responses that indicate when and how a prediction is incorrect.

Predictive processing has long been proposed as a core cortical function (Bastos et al., 2012; Keller and Mrsic-Flogel, 2018; Rao and Ballard, 1999). In many models, layer 2/3 pyramidal neurons compute prediction errors by comparing feedforward sensory-driven input with learned predictions. Layer 2/3 pyramidal neurons exhibit spiking responses resembling prediction errors across multiple modalities and paradigms (Audette and Schneider, 2023; Cole et al., 2024; Fiser et al., 2016; Furutachi et al., 2024; Hamm et al., 2021; Schneider et al., 2018; Solyga and Keller, 2025). Such responses have been studied particularly extensively in mouse V1 in the context of visuomotor prediction, where visual flow feedback in a virtual reality tunnel is coupled to locomotion speed. Layer 2/3 neurons display responses to perturbations in the expected visual motion during locomotion (Keller et al., 2012; Zmarz and Keller, 2016) and integrate visual flow and locomotion-related inputs with opposing signs (Attinger et al., 2017; Jordan and Keller, 2020) – a signature of comparative visuomotor prediction error computation. While this opposing integration depends on visuomotor experience and on local synaptic plasticity (Attinger et al., 2017; Widmer et al., 2022), how locomotion-related inputs and visual flow inputs become tuned to oppose each other in individual neurons remains unclear.

An answer to this could lie in the central learning mechanism of predictive processing: prediction errors drive corrective updates to improve the match, and thus cancellation, between predictive and sensory inputs – a process termed prediction error minimization. One candidate mechanism for this is that prediction errors, reflected in layer 2/3 firing, strengthen inhibitory inputs to better balance coincident excitation (Hertäg and Sprekeler, 2020). This process should progressively suppress neuronal responses to predictable excitatory input across experience – a phenomenon reported in layer 2/3 activity and driven by the increased inhibitory influence of top-down cortical inputs (Garner and Keller, 2022; Schneider et al., 2018; Tsukano et al., 2026). However, direct evidence that activity-dependent plasticity shapes this suppression is limited, leaving an important gap in the support for prediction error minimization, and, by extension, cortical predictive processing as a framework.

In addition to layer 2/3 neurons, neuromodulatory inputs from the locus coeruleus (LC) signal prediction errors to the cortex, including visuomotor errors (Basu et al., 2024; Jordan, 2024; Jordan and Keller, 2023; Su and Cohen, 2022). Because LC inputs convey a wide variety of different signals, their activity is likely uninformative of the specific prediction errors that occurred and thus cannot be used to directly instruct corrections to cortical predictions. Instead, the LC has been proposed to control the rate of predictive learning: as cortical prediction errors become more frequent, they elicit greater LC output to accelerate prediction error minimization (Jordan, 2024; Sales et al., 2019; Yu and Dayan, 2005). However, it remains unclear to what extent cortical activity shapes LC output, and although noradrenergic input facilitates several forms of cortical plasticity (Bear and Singer, 1986; Jordan and Keller, 2023; Martins and Froemke, 2015), whether this function extends to plasticity consistent with cortical prediction error minimization is unknown, in part because such plasticity is yet to be demonstrated.

Here, we test experimental predictions of prediction error minimization by focusing on visuomotor integration in layer 2/3 of mouse V1. We posit that spiking in layer 2/3 neurons during self-generated visual flow should drive a firing-rate-dependent enhancement of locomotion-related inhibition to suppress self-generated activity (**Figure 1A**). We test this using closed-loop manipulation of firing rates in individual layer 2/3 neurons and by examining calcium activity driven by self-generated visuomotor input. We find that spiking during self-generated visual flow drives changes in locomotion-related inputs with a direction dependent on the visual responsivity of the neuron. This results in more opposing visuomotor inputs, alongside reduced self-generated responses, consistent with an improvement of predictive visuomotor cancellation. We further show that LC input acutely facilitates this plasticity, and that cortical activity itself helps recruit arousal-linked neuromodulation, revealing an interaction between the cortex and subcortical neuromodulatory systems in the control of predictive visuomotor plasticity.

**Figure 1.**
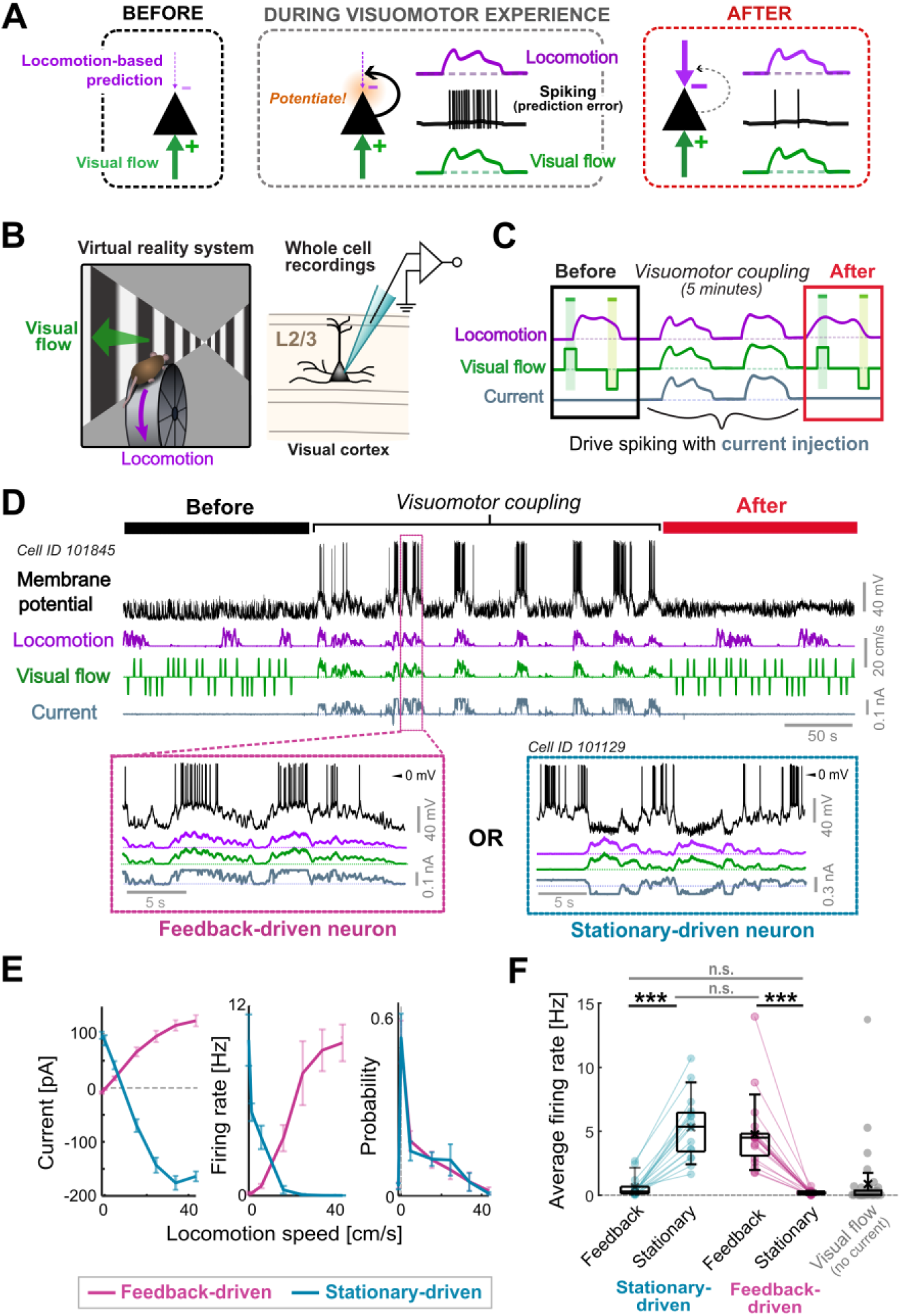
Whole cell recording experiment to test spiking-driven visuomotor plasticity. **(A)** Hypothesized prediction error minimization in V1 layer 2/3 neurons. Before visuomotor experience, visually driven excitation exceeds the locomotion-related inhibition. This imbalance leads to high firing rates during visuomotor feedback – i.e., a large prediction error, driving an enhancement of locomotion-related inhibition and reduced prediction error. The low firing afterwards prevents further updates to synaptic weights. **(B)** Schematic of the whole cell recording experiment. **(C)** Schematic of the virtual reality paradigm. 800 ms fixed-speed visual flow stimuli are presented before and after a visuomotor coupling period. Positive and negative deflections in visual flow indicate nasotemporal and temporonasal directions respectively. During visuomotor coupling, spiking is controlled with current injection coupled to locomotion. **(D)** Example trace from a neuron that underwent feedback-driven current injection. Bottom left shows an expanded section of this trace. Bottom right shows traces from a different neuron that underwent stationary-driven current injection to drive spiking when stationary. **(E)** From left to right: Average current injected, firing rate, and probability (i.e., fraction of time spent) as function of locomotion speed during the visuomotor coupling period, for stationary- and feedback-driven neurons. Error bars show SEM. **(F)** Average firing rates driven during feedback and stationary epochs of the visuomotor coupling period, compared for feedback-driven and stationary-driven neurons. Asterisks indicate the outcome of comparative tests: ***, p <0.001; n.s., p > 0.05. See **Statistical table S1** for details. Peak firing rate responses for all neurons during visual flow stimuli and locomotion (with no current injection) are shown in grey for a comparison to physiological firing rates.

## Results

We first asked whether neuronal spiking during experience of a predictable visuomotor relationship causally shapes the visuomotor inputs of layer 2/3 neurons in V1. Our hypothesis was that spiking coincident with visuomotor experience should promote greater predictive cancellation, primarily via the enhancement of locomotion-driven inhibition (**Figure 1A**).

To test this, we made whole cell current clamp recordings from 37 putative pyramidal neurons in layer 2/3 while mice ran head-fixed in a striped virtual reality tunnel (**Figure 1B**). Recordings began with a period in which locomotion produced no visual flow feedback; instead, brief fixed-speed visual flow stimuli were presented (**Figure 1C**), allowing us to measure both visual flow and locomotion-related influences on the membrane potential (V_m_). This was followed by a 5-minute period of visuomotor coupling, during which visual flow scaled with locomotion speed (‘visuomotor feedback’), providing experience of a predictive visuomotor relationship (**Figure 1C**). Afterwards, we repeated the fixed-speed visual flow stimuli to assess how visual and locomotion-related responses had changed.

To test the impact of spiking on changes in visuomotor input, we injected current to control firing rates during the visuomotor coupling experience. In 19 neurons, we injected depolarizing current proportional to locomotion speed to drive spiking during visuomotor feedback (‘feedback-driven’ neurons; **Figure 1D-E**). Assuming firing rates in layer 2/3 neurons reflect prediction errors, this simulates a persistent prediction error during self-generated visual flow (as depicted in **Figure 1A**). For comparison, we did the reverse in another 18 neurons: tonic depolarizing current evoked spiking when the mouse was stationary, while hyperpolarizing current suppressed spiking during visuomotor feedback (‘stationary-driven’ neurons; **Figure 1D-E**). Critically, induced firing rates for both groups were comparable, and were within the physiological range evoked by visual flow stimuli (**Figure 1F**). Firing rates were primarily determined by the current injected (which varied in gain across neurons) rather than average locomotion speeds, visuomotor response properties, or electrophysiological properties (**Figure S1**). This means that any relationship found between evoked firing rates and response changes is not confounded by these other factors.

### Spiking during visuomotor feedback drives changes in locomotion-related input

We first tested the hypothesis that spiking during visuomotor experience drives an enhancement of predictive inhibition during locomotion (**Figure 1A**).

To achieve this, we quantified each neuron’s locomotion-related input before and after visuomotor experience, using cross-correlations between subthreshold V_m_ and locomotion speed (**Figure 2A**). Positive and negative correlations indicate depolarization and hyperpolarization as locomotion speed increases, respectively. We measured changes in locomotion-related input as the difference in average locomotion-V_m_ correlation between the periods before and after visuomotor coupling experience, finding a range of different changes across neurons (**Figure 2A**).

**Figure 2.**
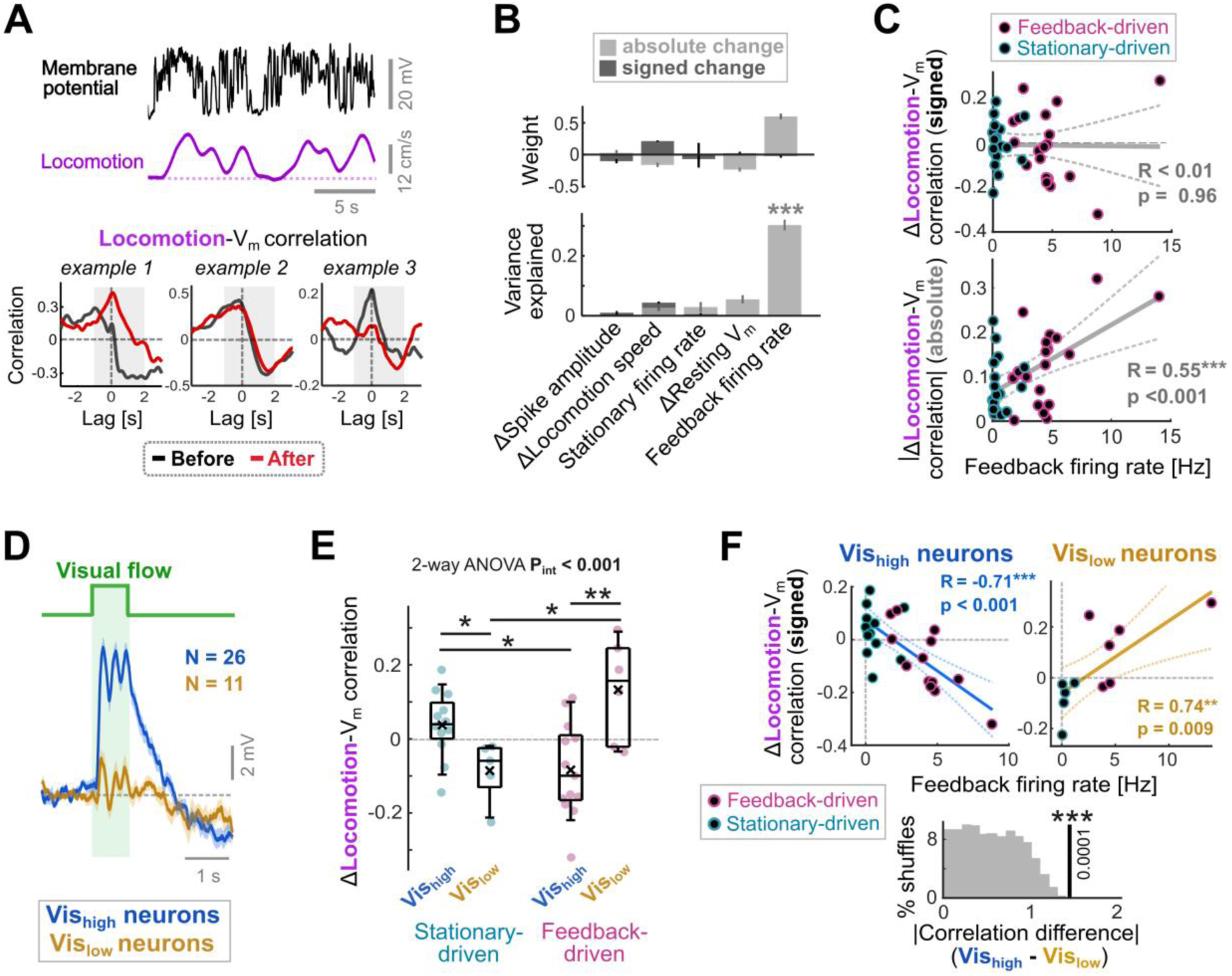
Changes in locomotion-related responses depend on firing rates during visuomotor feedback and visual responsivity. **(A)** Top: Membrane potential (V_m_) and locomotion speed from an example neuron. Bottom: Cross correlations computed between locomotion speed and V_m_ before (black) and after (red) the visuomotor coupling period for three example neurons. Shaded area shows time window used to compute average locomotion-V_m_ correlations. **(B)** Average weights and variance explained in the change in locomotion-V_m_ correlation for five different predictors, computed using multiple linear regression – see Methods. Error bars show SD, and asterisks indicate the p-value of the predictor; ***: p < 0.001. Light and dark gray indicates values for absolute and signed locomotion-V_m_ correlation changes respectively. **(C)** Change in average locomotion-V_m_ correlation (top: signed change, bottom: absolute change) plotted against firing rates during visuomotor feedback. **(D)** Average visual flow response across neurons with low visual flow responses (Vis_low_), and high visual flow responses (Vis_high_). Shading indicates SEM. **(E)** Locomotion-V_m_ correlation changes across groups defined by visual responsivity and current injection type. P_int_ refers to the p-value for the interaction between visual response type (Vis_high_ vs Vis_low_) and current injection type in a two-way ANOVA. Asterisks indicate p-values of comparisons: *: p < 0.05; **: p < 0.01. See **Statistical table S1** for details. **(F)** Change in locomotion-V_m_ correlation plotted against firing rates during feedback. Histogram shows distribution of differences in correlation between Vis_low_ and Vis_high_ neurons for 10000 shuffled datasets (grey distribution) and the actual data (black line). P-value < 0.001 (see Methods).

We next asked to what extent these changes were determined by the spiking we evoked during visuomotor experience compared to other factors. We assessed the predictive value of firing rates during visuomotor feedback and stationary periods, changes in locomotion speed, and correlates of recording stability (changes in baseline V_m_ and spike amplitudes), using multiple linear regression performed separately for signed and absolute changes in locomotion-V_m_ correlation. Firing rate during feedback emerged as the only significant predictor, explaining 30% of the variance in absolute locomotion-V_m_ correlation changes, while all other factors contributed negligible predictive value (**Figure 2B-C**). No factor, however, explained a significant share of variance in the signed changes in locomotion-V_m_ correlation.

Thus, high firing rates during visuomotor feedback led to greater changes in the influence on locomotion on the membrane potential, supporting the idea that locomotion-related input is altered in a firing rate-dependent manner during visuomotor experience. We did not, however, find evidence that firing consistently enhances locomotion-related inhibition across neurons.

### Visual flow input dictates the sign of locomotion-related response changes

Since firing rates during feedback could predict the magnitude of locomotion-related response changes, but not their direction, the latter must be governed by another factor. Functional groups in layer 2/3 are distinguishable by their visual flow, locomotion-related, and visuomotor mismatch responses, which correlate with differences in electrophysiological properties and molecular profiles (Jordan and Keller, 2020; O’Toole et al., 2023). These differential properties are consistent with these groups computing positive and negative prediction errors, reflecting unexpected excesses or omissions of expected visual flow, respectively. We hypothesized that neurons which receive visually driven excitation should be particularly subject to spiking-dependent enhancements of locomotion-related inhibition, since spiking during visuomotor feedback will be ‘interpreted’ by these neurons as an excess of unpredicted visually driven excitation (**Figure 1A**). If this is the case, spiking in visually excited neurons should primarily evoke negative changes in the influence of locomotion-related input on V_m_.

To test this, we grouped neurons by their responses to visual flow stimuli as Vis_high_ (mean depolarization >2 mV; n = 26) or Vis_low_ (n = 11; **Figure 2D**). A two-way ANOVA revealed a significant interaction between visual responsivity and current injection on signed changes in locomotion-V_m_ correlation, with feedback-driven Vis_high_ neurons alone showing a significant negative change (**Figure 2E**). Strikingly, Vis_high_ and Vis_low_ neurons both displayed significant relationships between their changes in locomotion-V_m_ correlation and firing rates during visuomotor feedback, but these relationships were in opposing directions: negative in Vis_high_ neurons, but positive in Vis_low_ neurons (**Figure 2F**). A permutation test confirmed that visual responsivity had a significant effect on these relationships (see Methods; **Figure 2F**). Notably, in a separate group of neurons driven to fire during locomotion in absence of visual flow feedback, no systematic changes in locomotion-V_m_ correlation were observed, suggesting that visual flow input is itself required for plasticity in locomotion-related input (**Figure S2**).

Together, these results show that in neurons with strong visually driven excitation, the changes in locomotion-related responses aligned with our hypothesis of an activity-dependent increase in predictive inhibition (**Figure 1A**). This was not, however, the case for neurons with weak or hyperpolarizing visual response. Since it is possible that the latter neurons may instead undergo enhanced visually-driven inhibition as a prediction error minimization mechanism (Attinger et al., 2017; Hertäg and Sprekeler, 2020), we performed similar analyses for changes in visual flow responses (**Figure S3**). These indicated that, while visual flow responses tended to reduce over time across neurons, such changes were not predictable from spiking evoked during visuomotor experience, or on visual responsivity. Thus, spiking may exert a more overt influence on updating predictive, motor-related inputs than sensory-driven inputs.

### Spiking during feedback drives the emergence of opposing visuomotor inputs

In mice highly trained with visuomotor coupling in VR, neurons in layer 2/3 of V1 integrate visual flow and locomotion-related inputs with opposing signs, a feature consistent with predictive visuomotor cancellation and prediction error computation (Attinger et al., 2017; Jordan and Keller, 2020; Widmer et al., 2022). If prediction error minimization underlies this organization, then firing during visuomotor feedback should enhance the opposing relationship between visuomotor inputs.

To assess this, we first compared locomotion-related responses across groups defined by visual flow responsivity and current injection type. A two-way ANOVA revealed no significant differences in locomotion-V_m_ correlations between Vis_high_ and Vis_low_ neurons prior to visuomotor coupling (**Figure 3A**). After visuomotor coupling, however, significant differences emerged: In feedback-driven neurons, locomotion-V_m_ correlations became significantly lower in Vis_high_ compared to Vis_low_ neurons (consistent with an opposing visuomotor input organization), and both groups showed significant differences relative to their stationary-driven counterparts (**Figure 3A**).

**Figure 3.**
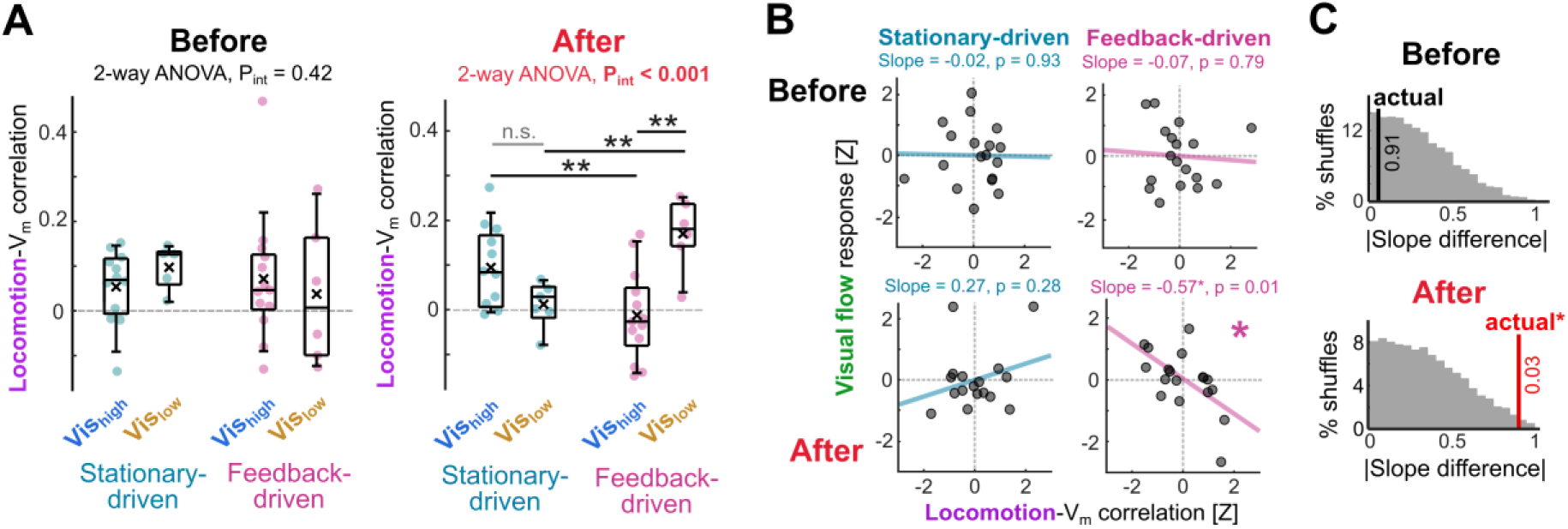
Spiking during visuomotor feedback generates stronger opposition between visual and locomotion-related responses. **(A)** Locomotion-V_m_ correlations, before and after visuomotor coupling, across groups defined by visual responsivity and current injection type. P_int_ refers to the p-value for the interaction between visual response type and current injection type in a two-way ANOVA; all other ANOVA p-values exceeded 0.05. Asterisks indicate p-values of pairwise comparisons: *: p < 0.05; **: p < 0.01. See **Statistical table S1** for details. **(B)** Visual flow responses plotted against locomotion-V_m_ correlations (z-scored) before and after the visuomotor coupling period, for feedback-driven and stationary-driven neurons. Slope of linear model fit is indicated – i.e., the ‘visuomotor slope’. **(C)** Histograms of absolute difference in visuomotor slope between stationary-driven and feedback-driven neurons for 10000 datasets permuted with respect to experimental group (see Methods). Actual difference is indicated with a bold line, with p-values shown.

We next quantified this visuomotor input organization directly, as the slope of the linear fit between visual flow and locomotion-related responses across neurons (the ‘visuomotor slope’). Before visuomotor coupling, both feedback-driven and stationary-driven neurons displayed little relationship between visual and locomotion-related responses (**Figure 3B**), and their visuomotor slopes did not differ significantly. After visuomotor coupling, however, feedback-driven neurons developed a significant negative relationship between visual and locomotion-related responses, whereas stationary-driven neurons still displayed little relationship between the two, resulting in a significant difference emerging between the visuomotor slopes for feedback-driven and stationary-driven neurons (**Figure 3C**).

Together, these results indicate that spiking during feedback shapes visuomotor integration such that locomotion-related inputs and visual flow inputs become more opposing – a signature of enhanced visuomotor prediction error computation.

### Activity-dependent reorganization of visuomotor integration is evident in population calcium activity

The spiking-driven response changes described so far were quantified from subthreshold activity due to the sparse firing of layer 2/3 neurons, while sample size was limited owing to the low throughput of whole cell recordings in vivo. We therefore assessed whether similar plasticity was evident at the population output level by analyzing a published two-photon calcium imaging dataset (Jordan and Keller, 2023). Here, jGCaMP8m fluorescence was recorded from V1 layer 2/3 neurons while mice experienced the same sequence of visual flow stimuli and visuomotor coupling as in the whole cell recordings (**Figure 4A-B**). In the dataset analyzed first, noradrenergic locus coeruleus (LC) axons in the imaging site were periodically optogenetically stimulated during the visuomotor coupling period. This manipulation was previously shown to facilitate plasticity resulting in reduced responses to visual flow stimuli during locomotion (Jordan and Keller, 2023). However, the activity dependence of plasticity, changes in locomotion-related responses, and changes in visuomotor integration were not previously examined. We therefore tested this dataset for the two major activity-dependent effects observed in the electrophysiology experiments: 1) Locomotion-related response changes whose sign depend on visual responsivity (**Figure 2**), and 2) Enhanced opposition between visual and locomotion-related inputs (**Figure 3**).

**Figure 4.**
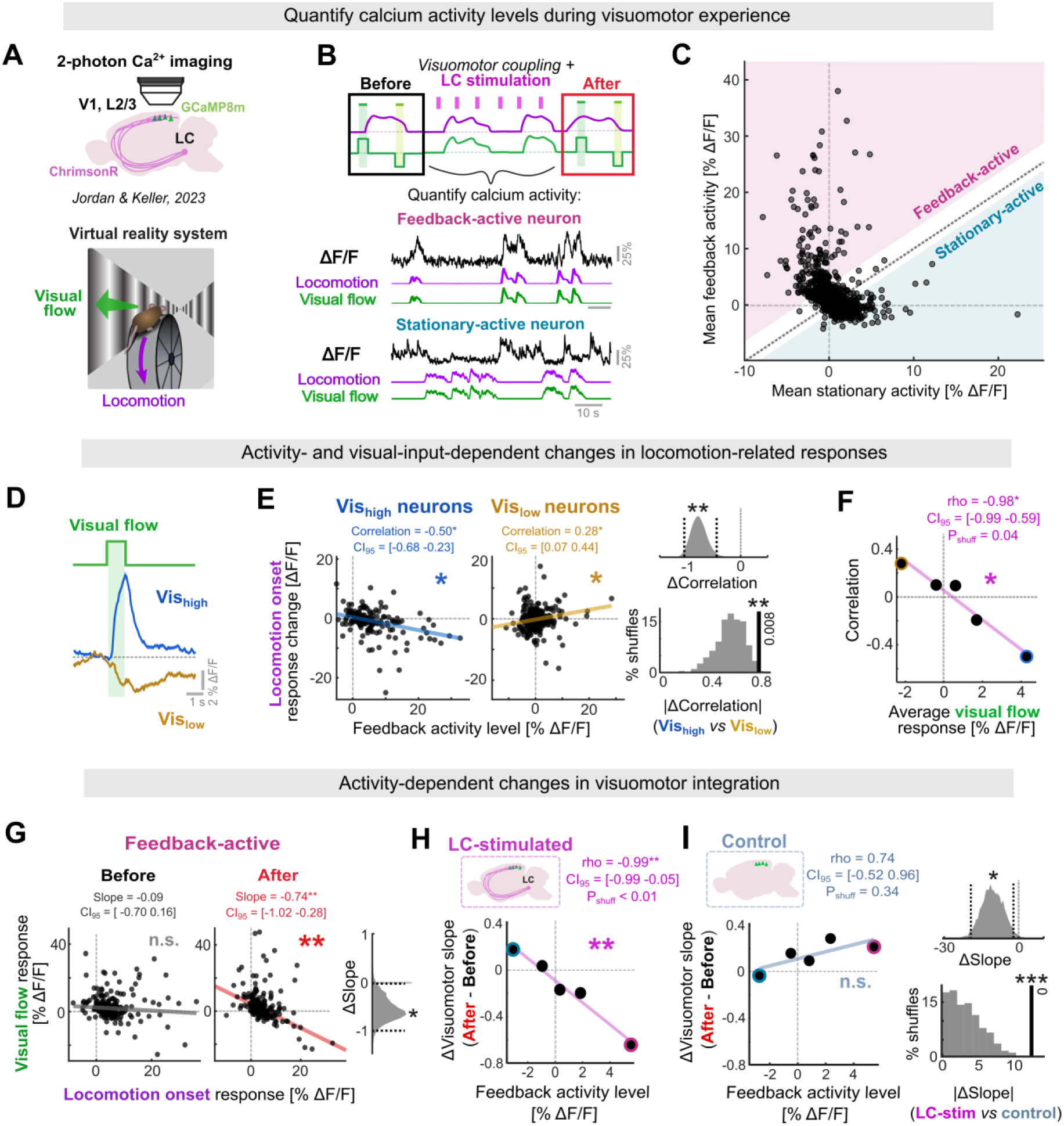
Activity-dependent changes in visuomotor input in population calcium activity. **(A)** Schematic of two-photon imaging in layer 2/3 of V1 during virtual reality experience - dataset from (Jordan and Keller, 2023). In the dataset analyzed in panels A–H, LC axons were densely labelled with ChrimsonR. **(B)** Sequence of stimuli. Visual flow stimuli were presented before and after 5- or 10-min visuomotor coupling with optogenetic stimulation of LC axons. Example calcium traces from a feedback-active and a stationary-active neuron are shown. **(C)** Mean calcium activity during visuomotor feedback versus stationary periods across all neurons during visuomotor coupling. Shaded regions indicate thresholds used to classify feedback-active and stationary-active neurons. **(D)** Mean calcium responses to visual flow stimuli in Vis_high_ and Vis_low_ neurons. Shading indicates SEM. **(E)** Locomotion onset response changes (after − before visuomotor coupling) versus feedback activity level for Vis_high_ and Vis_low_ neurons. 95% confidence intervals (CI_95_) and p-values were obtained by hierarchical bootstrapping (see Methods). Violin plot shows the bootstrapped distribution of correlation differences between Vis_high_ and Vis_low_ neurons, with CI_95_ indicated by black dotted lines. Histogram compares absolute slope differences with those from 1000 datasets permuted with respect to visual flow responses (see Methods). **(F)** Correlation coefficients between feedback activity level and locomotion onset response change (as in panel E) plotted against median visual flow responses for quintiles of the data selected by visual flow responses. P_shuff_ was derived from permutation analysis; CI_95_ values were obtained by hierarchical bootstrapping (see Methods). **(G)** Relationship between visual flow and locomotion onset responses before and after visuomotor coupling for feedback-active neurons. CI_95_ and p-values for linear fit slopes (visuomotor slopes) were obtained by hierarchical bootstrapping (see Methods). Violin plot shows the distribution of slope changes (after – before) across 1000 hierarchically bootstrapped datasets; black dotted lines indicate CI_95_. **(H)** Change in visuomotor slope (as in panel G) plotted against median feedback activity level for five dataset quintiles grouped by feedback activity level. P_shuff_ was derived from permutation analysis; CI_95_ values were obtained by hierarchical bootstrapping (see Methods). **(I)** As in panel H, but for a control dataset without ChrimsonR in LC axons. Violin plot shows the hierarchically bootstrapped distribution of slope differences between control and LC-stimulated datasets (black dotted lines indicate CI_95_). Histogram below compares absolute actual slope difference with those from 1000 datasets permuted with respect to experimental group (see Methods).

We began by quantifying each neuron’s level of calcium activity during visuomotor feedback (‘feedback activity level’), calculated as the difference in average activity during visuomotor feedback relative to stationary behavior of the visuomotor coupling period (**Figure 4B-C**). Neurons spanned a range from strongly activated to suppressed during feedback, mirroring the firing patterns we evoked via current injection in the whole-cell recordings.

We then tested whether these patterns of calcium activity predicted changes in locomotion-related input across visuomotor experience, with a direction of change set by visual responsivity – as observed in the electrophysiology (**Figure 2**). To measure changes in locomotion-related inputs, we quantified calcium responses to locomotion onsets before and after visuomotor coupling experience, while responses to visual flow stimuli in the same periods were used to group neurons by visual responsivity (**Figure 4D**). In Vis_high_ neurons (top 20% of visual flow responses) changes in locomotion onset responses displayed a significant negative correlation with feedback activity levels: i.e., reductions in locomotion onset responses were associated with higher calcium activity during visuomotor feedback (**Figure 4E**). Vis_low_ neurons (lowest 20% of visual flow responses) displayed the opposite pattern: a positive relationship that significantly differed from that of the Vis_high_ neurons (**Figure 4E**). This divergence was systematic: across subsets of the data grouped by visual responsivity, the correlation between feedback activity levels and changes in locomotion onset response became progressively more negative as the average visual flow response increased (**Figure 4F**) – a relationship confirmed significant by both hierarchical bootstrapping and permutation tests (see Methods). Thus, changes in locomotion-related responses were predictable from activity levels during visuomotor experience, while visual responsivity dictated the sign of change, mirroring the effect observed in the whole cell recordings (**Figure 2**). In contrast, we did not find evidence for activity-dependent changes in visual flow responses that were predictable from locomotion responsivity (**Figure S4**).

We next tested whether visuomotor inputs became more opposing dependent on feedback activity levels, as observed in the whole cell recordings (**Figure 3**). Relationships between visual flow and locomotion onset responses were measured from the periods before and after visuomotor coupling. In feedback-active neurons (with feedback activity levels in the top 20%; **Figure 4C**), the opposing relationship between these responses was significantly enhanced after visuomotor coupling, evident in the slope of the linear fit (i.e., ‘visuomotor slope’) (**Figure 4G**), analogous to the changes seen in feedback-driven neurons of the whole cell recordings. This effect scaled with the level of calcium activity during visuomotor feedback: subsets of neurons with higher average feedback activity levels showed correspondingly larger negative shifts in visuomotor slope following coupling (**Figure 4H**) – a significant relationship according to both hierarchical bootstrapping and permutation tests.

Thus, both signatures of activity-dependent visuomotor plasticity identified in the whole-cell recordings had clear parallels at the level of population calcium activity. This convergence suggests that visuomotor-driven neuronal activity can elicit plasticity consistent with prediction error minimization, and that this plasticity is detectable at the level of neuronal output.

### LC axonal stimulation facilitates activity-dependent visuomotor reorganization in calcium activity

In the calcium imaging dataset analyzed so far, LC axons within the imaging window were optogenetically stimulated at intervals during visuomotor coupling experience. The LC has been proposed to control the rate of predictive cortical learning (Jordan, 2024; Sales et al., 2019; Yu and Dayan, 2005). If this is the case, then activity-dependent visuomotor plasticity should be weaker across the same timescale when LC input is not artificially elevated.

To test this, we compared the activity-dependent changes in visuomotor responses observed in the LC-stimulated dataset to those in a control dataset, in which ChrimsonR expression was absent or negligible in cortical-projecting LC axons (**Figure 4I**) (Jordan and Keller, 2023). Feedback activity levels and locomotion behavior during visuomotor coupling were comparable between LC-stimulated and control datasets, indicating no gross impact of LC stimulation on calcium activity levels or on visuomotor experience (**Figure S5A-B**), consistent with previous conclusions (Duss et al., 2026; Jordan and Keller, 2023). Strikingly, however, the activity-dependent enhancement of opposing visuomotor input observed in LC-stimulated mice (**Figure 4H**) was not evident in control mice (**Figure 4I and S5C**). The relationship between feedback activity levels and changes in visuomotor slope differed significantly between LC-stimulated and control datasets, as determined by both permutation and hierarchical bootstrap tests (**Figure 4I** – see Methods). The same was true of activity-dependent changes in locomotion-related responses, which were also not evident in control mice, and displayed little relationship with visual responsivity (**Figure S5D-F**).

Together, this indicates that optogenetic elevation of LC output during visuomotor experience facilitates activity-dependent reorganization of visuomotor input in layer 2/3, in line with the hypothesized role of the LC as a rate-controller of cortical predictive learning.

### Activity-dependent reduction in self-generated activity parallels development of opposing visuomotor inputs

Prediction error minimization results in reductions in predictable neuronal activity across experience, via the increased cancellation between sensory and predictive inputs (**Figure 1A**). If the activity-dependent reorganization of visuomotor inputs described so far reflects this process, then reductions in predictable, visuomotor-driven neuronal activity should occur in parallel. Therefore, we next tested for activity-dependent reductions in self-generated activity across visuomotor coupling experience in the calcium imaging dataset.

To measure changes in self-generated activity, we quantified how neuronal responses to feedback onsets (initiations of locomotion and visual flow feedback) changed between the beginning of the visuomotor coupling period (early), and the end (late) (**Figure 5A**). In mice with stimulation of LC axons, feedback-active neurons displayed a significant reduction in feedback onset responses across visuomotor experience (**Figure 5A**) that could not be explained by changes in locomotion behavior (**Figure S6A**). Across the dataset and in individual imaging sites, reductions in feedback onset responses were strongly dependent on calcium activity levels during visuomotor feedback (**Figure 5B and S6B**) and paralleled the negative changes in visuomotor slope across subsets of the data (**Figure 5C**). These results are in line with the activity-dependent enhancement of the opposing organization between visuomotor inputs reflecting an improvement in predictive visuomotor cancellation, leading to reduced self-generated activity (**Figure 1A**).

**Figure 5.**
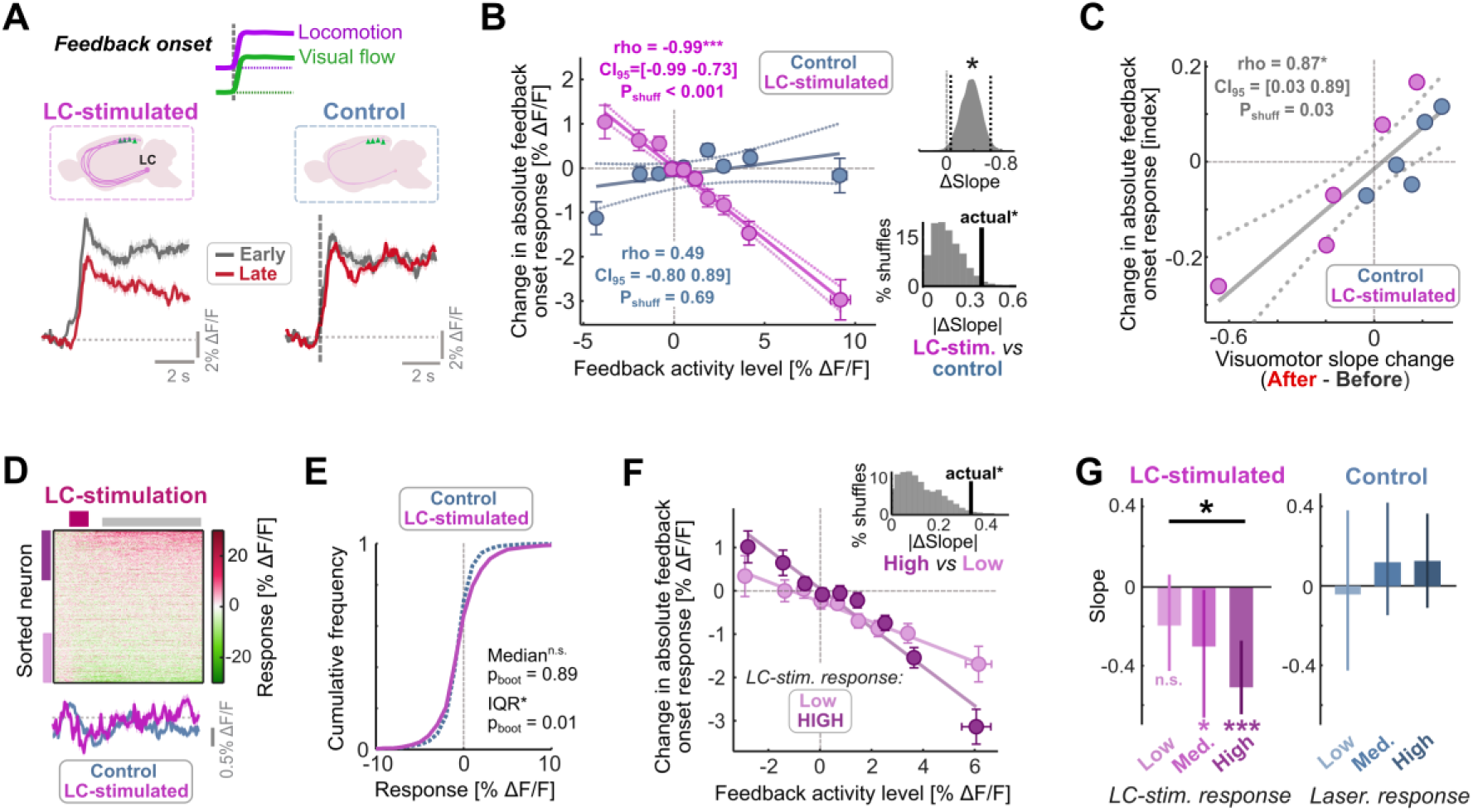
LC axonal stimulation facilitates activity-dependent reductions in self-generated calcium activity. **(A)** Calcium responses to feedback onsets averaged across feedback-active neurons early (black) and late (red) in the visuomotor coupling period. Shaded area indicates SEM. Responses are shown for mice undergoing optogenetic stimulation of LC axons during visuomotor experience (LC-stimulated), and control mice lacking opsin expression. **(B)** Left: Average change in the absolute feedback onset response plotted against average feedback activity level for data subsets grouped by feedback activity levels (see Methods). Error bars indicate SEM. Violin plot shows distribution of slope differences between control and LC stimulated bootstrapped datasets (confidence intervals shown in dotted black), while below histogram compares the absolute difference in linear fit slopes with those of 1000 permuted datasets – see Methods. **(C)** Average change index in absolute feedback onset response (a measure of change normalized according to response sizes – see Methods) plotted against the change in visuomotor slope (as shown in Figure 4G-I) for 20% subsets of the data grouped by feedback activity levels. P-value is derived from a shuffle analysis. **(D)** Heatmap of calcium responses to optogenetic laser stimulation across all LC-stimulated neurons, sorted by average response. Gray bar indicates window used to calculate average responses. Plot below shows average across all LC-stimulated neurons and control neurons. Shading indicates SEM. **(E)** Cumulative histogram comparing average responses to optogenetic laser stimulation in LC-stimulated and control groups. P values (P_boot_) are shown for outcome of hierarchical bootstrap tests for differences in median and inter-quartile range (IQR) between LC-stimulated and control groups – **see Methods and Statistical table 2**. **(F)** Similar to panel B, but where neurons are split into those with a high average response to LC axon stimulation (top third) and those with a low response (bottom third) within each group. Histogram compares the actual difference in linear fit slopes with those of 1000 datasets permuted for LC stimulation response – see Methods. **(G)** Slope of the fits shown in panel F compared for neurons with high, low, and intermediate (‘Med.’) responses to LC axon stimulation (left) and optogenetic laser stimulation in control mice (right). Error bars indicate 95% confidence intervals computed using hierarchical bootstrapping. Asterisks indicate the P_boot_ values resulting from hierarchical bootstrapping: *: P_boot_ < 0.05; ***: P_boot_ < 0.001.

Activity-dependent reductions in self-generated activity were notably absent in control animals lacking ChrimsonR in LC axons, where the relationship between feedback activity levels and changes in feedback onset response was significantly more positive than that in LC-stimulated mice (**Figure 5A-B**). This reinforced the idea that plasticity consistent with prediction-error minimization is acutely facilitated by elevated LC input during visuomotor experience.

### Calcium activity following early LC stimulation predicts the strength of plasticity

One way LC output could enhance neuronal plasticity, and thus prediction error minimization, is by enhancing evoked layer 2/3 calcium activity – for example, via effects of noradrenaline on voltage gated calcium channels (Gray et al., 2021; Marzo et al., 2009). LC stimulation evoked no systematic, immediate calcium response (Jordan and Keller, 2023), and had no gross effect feedback activity levels in layer 2/3 neurons (**Figure S5A**), but these analyses did not rule out slow, transient, or subtle effects on evoked calcium activity. We therefore examined in more detail whether LC axonal stimulation affected calcium activity during visuomotor feedback, and whether any such effect predicted the magnitude of subsequent response changes.

Given the slow dynamics of cortical noradrenaline (Matarasso et al., 2025; Neyhart et al., 2026), we quantified calcium responses 2-8 s after LC stimulation during visuomotor feedback early in the coupling period. While there was no systematic response on average across neurons (**Figure 5D**), calcium activity changes following LC stimulation displayed significantly higher variance than those evoked by the optogenetic laser in control mice (**Figure 5E**). One possibility is that this heterogeneity reflects the proposed role of the LC as a cortical gain-enhancer (Aston-Jones and Cohen, 2005), in which case, stimulating the LC during feedback should result in response changes that scale with the feedback activity level of the neuron. However, LC stimulation responses displayed a mild but non-significant dependence on feedback activity levels (**Figure S6C-D**), arguing against gain modulation underlying most of the response heterogeneity. We posit that it could instead be caused by heterogeneous opsin labelling of LC axons (such that some neurons are subject to greater noradrenaline increases following stimulation than others) or by diversity in adrenergic receptor expression.

We next asked whether the heterogeneous response profile reflects a neuromodulatory influence that facilitates plasticity and varies in strength across neurons. Supporting this interpretation, the strength of activity-dependent reductions in self-generated activity scaled with responses to LC axonal stimulation across neurons (**Figure 5F**): neurons in the top third of responsiveness displayed a stronger relationship between feedback activity levels and changes in feedback onset responses than neurons in the bottom third – a significant effect according to hierarchical bootstrapping and permutation tests, and absent in control mice (**Figure 5G**). This was unlikely to reflect chance or an analytical artifact: categorizing neurons in an identical way based on their responses to sham onsets, rather than LC-stimulation, rarely reproduced the same effect size (**Figure S6E**).

Together, these results show that the interaction between neuronal activity and LC-evoked calcium responses predicted the degree of change in self-generated activity across visuomotor experience. This supports the idea that LC-driven neuromodulation controls the gain of prediction-error-driven plasticity at the level of individual neurons.

### Activity in the visual cortex can evoke pupil-linked neuromodulation

Finally, we turned our attention to the reverse relationship: the effect of visual cortical activity on neuromodulatory activity. Cortical LC axons display activity consistent with visuomotor prediction error signaling, similar to V1 layer 2/3 neurons (Jordan and Keller, 2023). Cortical neurons may thus facilitate prediction error minimization by contributing to the recruitment of plasticity-promoting neuromodulatory feedback.

We tested this idea using pupil diameter as a proxy for cortical neuromodulatory input, given its established correlations with cholinergic, serotonergic, and particularly noradrenergic axonal activity in the cortex (Jordan and Keller, 2023; Reimer et al., 2016; Yogesh and Keller, 2025), and the fact that activity in cortical LC axons is causally linked to pupil dilation (Jordan and Keller, 2023). To validate this proxy for neuromodulatory responses consistent with visuomotor prediction errors, we examined the relationship between evoked pupil dilations and LC axonal calcium activity in the cortex by analyzing our previously published 2-photon imaging dataset (Jordan and Keller, 2023) (**Figure 6A**). During imaging, mice experienced visuomotor coupling interspersed with visuomotor mismatch stimuli (1 s pauses in the visual flow during feedback), used to elicit visuomotor prediction error responses. When mismatch events were split into quintiles by evoked pupil derivative, LC axonal responses scaled linearly with average pupil dilation (**Figure 6B**). LC responses also correlated well with pupillary responses on a trial-by-trial basis within individual sessions (**Figure 6C**). Pupil dilations were therefore a good predictor of the cortical LC axonal activity evoked by visuomotor mismatch events, and we proceeded to use pupil diameter as a proxy for cortical noradrenergic input. Note that the axons of other pupil-linked neuromodulatory centers, by contrast, have not been found to respond to visuomotor mismatches (Yogesh and Keller, 2025, 2024).

**Figure 6.**
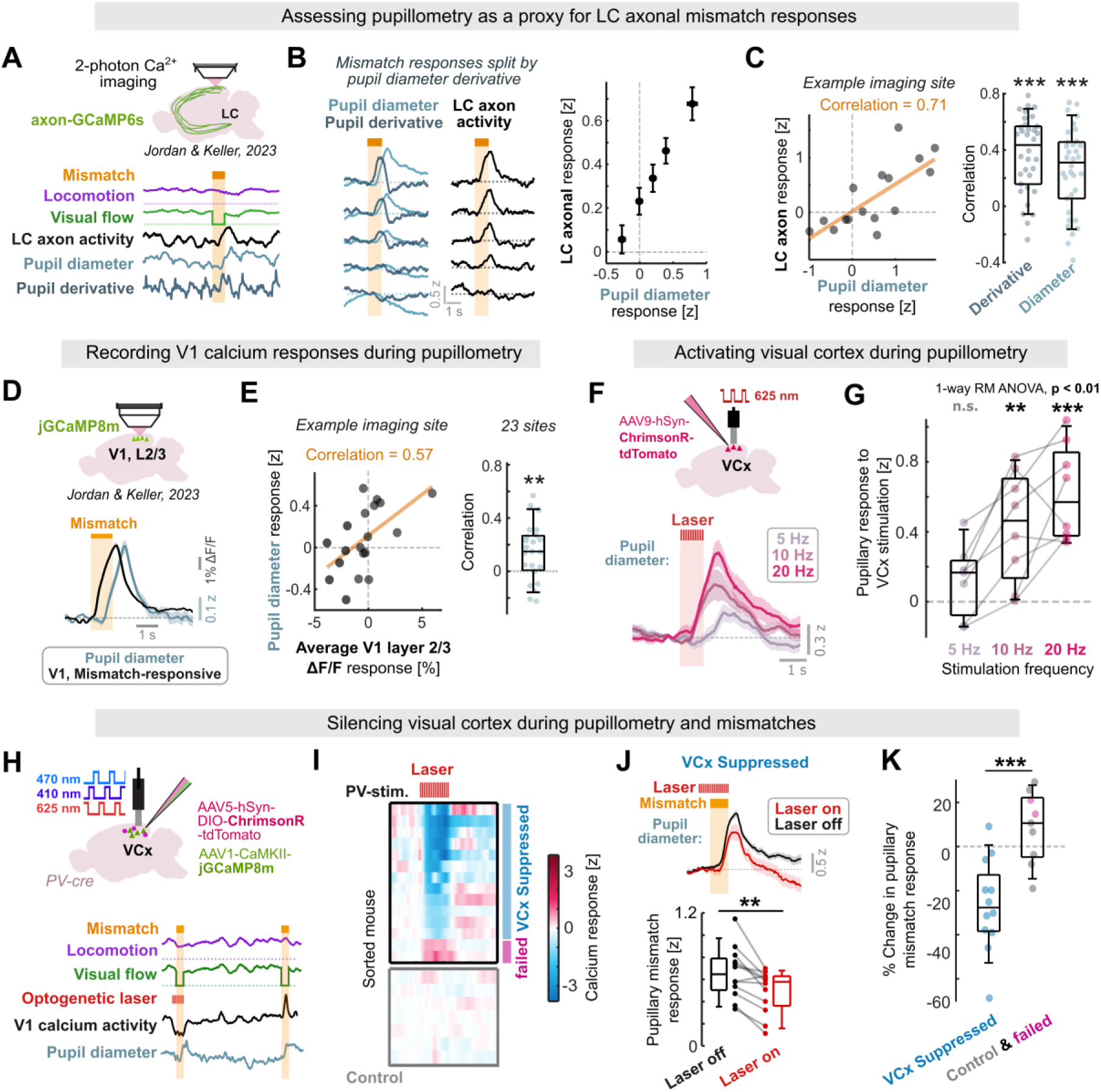
Visual cortex contributes to pupil-linked neuromodulation. **(A)** Example from 2-photon calcium imaging dataset (Jordan and Keller, 2023) in which LC axons were imaged in the cortex during visuomotor feedback and mismatch stimuli. **(B)** Left: Mismatch responses in pupil diameter, its time derivative, and LC axonal calcium activity averaged across trials split into quintiles by pupil derivative response. Trials were concatenated from across recording sites. Right: average pupil dilation plotted against the average LC axonal response for each data quintile. Error bars show SEM across trials. **(C)** Correlation between LC axonal calcium responses and pupil diameter responses to visuomotor mismatch stimuli across trials for an example imaging site. Boxplot shows respective correlation coefficients across all imaging sites for pupil diameter and pupil derivative responses. Asterisks indicate outcome of paired t-test (***: p < 0.001). **(D)** Calcium activity of layer 2/3 neurons in V1 was analyzed from a 2-photon imaging dataset (Jordan and Keller, 2023) in relation to concurrently recorded pupil diameter. Plots show average mismatch response across all neurons with an average response exceeding 1% ΔF/F, alongside the average pupillary response. Shading indicates SEM. **(E)** Correlation between V1 ΔF/F responses and pupil diameter responses to visuomotor mismatch across trials for an example imaging site. Boxplot shows correlation coefficients for all imaging sites. Asterisks indicate outcome of paired t-test (** p < 0.01). **(F)** Optogenetic activation of the visual cortex during pupillometry. Plots show the average pupil diameter responses to 5, 10, and 20 Hz stimulation across mice. **(G)** Average pupil responses to V1 optogenetic stimulation. Asterisks indicate outcome of paired t-test (n.s.: p > 0.05; ** p < 0.01; *** p < 0.001) (see **Statistical table S2** for details). **(H)** Schematic of simultaneous fiber photometry of pyramidal neuron calcium activity and silencing via optogenetic parvalbumin (PV) interneuron stimulation in the visual cortex. Example traces are shown below. On some visuomotor mismatch trials, PV neurons would be activated via an optogenetic laser to suppress visual cortex activity. Pupil diameter was recorded concurrently. **(I)** Heatmap of V1 calcium responses to optogenetic stimulations, in mice with stimulation of PV-interneurons (‘PV stim.’), and control mice lacking ChrimsonR expression. Blue bar indicates mice with significant activity suppression. **(J)** Average pupil dilation response to visuomotor mismatch with (red) and without (black) concurrent V1 suppression from an example mouse. Shading indicates SEM. Boxplot shows average pupil diameter responses to mismatch between trials with and without the optogenetic laser across all mice with significant V1 suppression. P-value indicates outcome of paired t-test (p < 0.01). **(K)** Changes in pupillary mismatch response between trials with the optogenetic laser on and off, plotted against the effect of the laser on V1 calcium activity for all mice.

To assess whether visuomotor mismatch responses in V1 neurons may contribute to neuromodulatory activation, we next analyzed datasets in which pupil diameter and V1 layer 2/3 calcium activity were concurrently measured during the presentation of visuomotor mismatches (**Figure 6D**). The average delay between layer 2/3 mismatch responses and pupil dilation was 595 ms (**Figure S7A-B**), while the amplitude of population responses to visuomotor mismatches in layer 2/3 correlated mildly but positively with pupillometric responses across trials in almost all imaging sites (**Figure 6E**), suggesting that LC activity could be influenced by the preceding V1 activity.

To test for a causal relationship, we next asked whether visual cortex activity can, in principle, recruit pupil-linked neuromodulation. We expressed ChrimsonR in visual cortical neurons and implanted an optic fiber above V1 to deliver 5-7 mW 635 nm excitation light while recording pupil diameter. Optogenetic stimulation of visual cortex neurons evoked pupil dilations with an amplitude that scaled with stimulation frequency (**Figure 6F**) and could not be explained by movement (**Figure S7C**). The onset time of pupil responses was 578 ms on average, similar to the delay between mismatch-evoked V1 and pupillary responses (**Figure S7B**).

Finally, to directly test if cortical visuomotor prediction-error-like signals contribute to pupillary-responses, we performed optogenetic silencing of the visual cortex: ChrimsonR was expressed in parvalbumin (PV) positive interneurons via AAV injections in PV-Cre mice (Hippenmeyer et al., 2005), while jGCaMP8m was expressed in pyramidal neurons under the CaMKIIa promoter (**Figure 6H**). In control mice, ChrimsonR expression was omitted. Using fiber photometry, we confirmed that activating PV interneurons suppressed pyramidal neuron calcium activity in all but two mice (which lacked ChrimsonR-tdTomato expression in post-hoc histology) (**Figure 6I**). In mice where cortical activity was successfully suppressed, pupillary mismatch responses were significantly reduced during optogenetic stimulation (**Figure 6J**), and the changes were significantly different from those in mice lacking cortical suppression, where no systematic effect of stimulation was observed on the pupillary response (**Figures 6K and S7D**). Thus, the reductions in pupillary response could not be explained by optogenetic excitation light evoking the pupillary light reflex, nor could they be explained by changes in locomotion speed (**Figure S7E**).

Together, these results show that visual cortical activity contributes to pupil-linked neuromodulatory responses that are consistent with visuomotor prediction errors. Given that pupillary responses are a good predictor of LC axonal activity in cortex, this supports a role for cortical activity in recruiting plasticity-promoting neuromodulatory feedback to facilitate prediction error minimization.

## Discussion

Predictive processing models propose that the cortex uses prediction error signals to learn and update predictions of the sensory world (Bastos et al., 2012; Keller and Mrsic-Flogel, 2018; Rao and Ballard, 1999). While responses consistent with prediction errors are well documented in layer 2/3, evidence for activity-dependent plasticity consistent with prediction error minimization has been lacking. Our results address this gap, providing evidence for plasticity in layer 2/3 of V1 consistent with prediction error minimization in the visuomotor domain: prediction errors, represented in layer 2/3 neuronal firing rates, drive changes that establish opposing locomotion-related and visual inputs, enhancing their cancellation to reduce the predictable components of activity. LC stimulation facilitates this plasticity, which could function to enhance learning rates when predictions are highly inaccurate – i.e., when the cortex signals a high rate of errors and drives high LC activity (Jordan, 2024). Consistent with this idea, our results demonstrate that visual cortex activity can recruit pupil-linked neuromodulation, suggesting that cortical spiking may mediate prediction error minimization in parallel ways: directly, through local plasticity, and – when population activity sufficiently high – by recruiting plasticity-promoting neuromodulatory systems like the LC.

Cortical predictions are, in principle, determined by both the activity of predictive inputs and their synaptic weights onto pyramidal neurons. Our results indicate that locomotion-related inputs can be adjusted for each neuron based on postsynaptic activity, with the sign of change determined by the visual responsivity of the neuron (**Figures 2 and 4D-F**). This effect could result from molecular (O’Toole et al., 2023) or circuit-level (Leinweber et al., 2017) differences correlating with visual responsivity, or from a direct influence of visually-driven postsynaptic activity itself – an interpretation supported by the absence of activity-dependent changes in locomotion-related input when visual flow feedback was omitted (**Figure S2**). If Vis_high_ neurons receive locomotion-related input predominantly on their basal dendrites (colocalizing with feedforward visual excitation) while Vis_low_ neurons receive locomotion-related input on their apical dendrites (consistent with a top-down input), then differential plasticity rules in different dendritic compartments offer another possible mechanism (Mikulasch et al., 2023; Rajan et al., 2026; Wright et al., 2025). This would align with previously postulated circuitry for positive and negative prediction error neurons (Keller and Mrsic-Flogel, 2018).

The negative change in locomotion-related responses in Vis_high_ neurons fits a prediction error minimization account: firing during feedback should increase locomotion-driven inhibition to cancel predictable activity (**Figure 1A**). The opposite sign of change in Vis_low_ neurons is not expected from prediction error minimization, which presumably must occur via another mechanism, such as enhancement of visually driven inhibition (Hertäg and Sprekeler, 2020). We found little evidence for spiking-dependent changes in visual input here (**Figures S3 and S4**), despite prior reports of such plasticity following visuomotor coupling experience (Attinger et al., 2017). It is possible that enhancement of visually driven inhibition depends on a measure of neuronal activity (e.g., dendritic voltage) or neuronal properties not captured in our recordings, or is only visible during precise matching of visual and motor inputs (and thus is not readily quantifiable during our open-loop conditions).

Some experimental predictions of prediction error minimization remain untested here. Top-down inputs (or the interneurons they target) are thought to undergo activity-and synapse-level adjustment to alter predictive input to layer 2/3. While cingulate afferents to V1 show activity consistent with a locomotion-based prediction (Leinweber et al., 2017), other sources of locomotion-related activity to V1 could also contribute to the changes we observe, such as thalamic inputs (Nestvogel and McCormick, 2022; Roth et al., 2016). Because V_m_ and calcium recordings cannot distinguish excitatory from inhibitory input, the reduced locomotion-related responses in Vis_high_ neurons, while consistent with increased inhibition, could equally reflect decreased excitation. Other plasticity principles may therefore account for the results, such as proposed mechanisms for causal inference (Mikulasch et al., 2022) or decorrelation of feedforward and feedback inputs (Rajan et al., 2026). Identifying the origin of activity-dependent changes in locomotion-related or other predictive inputs will be an important goal for constraining models of cortical function.

## Acknowledgements

We thank the Jordan lab for discussion, Bioresearch and Veterinary Services at the University of Edinburgh for animal husbandry, Nathalie Rochefort’s lab and Lida Zoupi’s lab for PV-cre mice, and Georg Keller, Nathalie Rochefort, Ian Duguid, Anna Vasilevskaya, Mahesh Karnani, and Loreen Hertäg for feedback on the manuscript. This work was supported by a grant from the Simons Foundation [AN-NC-GB-Independence Faculty-00872616, R.J.], supported by the Simons Initiative for the Developing Brain, and funded by the European Union [ERC, Learn2predict, 101162130, R.J.]. Views and opinions expressed are those of the authors only and do not necessarily reflect those of the European Union or the European Research Council. Neither the European Union nor the granting authority can be held responsible for them.

## Author contributions

S.Y.Y. and R.J. designed and performed experiments, M.B. performed surgery, behavioral training of animals, and histology, R.J. and S.Y.Y. conducted analyses, and R.J. and S.Y.Y. wrote the paper with input from all authors.

## Data and code availability

The whole cell recording dataset, optogenetics datasets, and analytical code will be made publicly available alongside peer-reviewed publication. The two-photon calcium imaging datasets were published previously (Jordan and Keller, 2023) and are publicly available (<u>10.5281/zenodo.8006971</u>).

## Declaration of interests

The authors declare no competing interests.

## Methods

### Animals and animal procedures

Animal experiments underwent review by the University of Edinburgh Animal Welfare and Ethical Review Body (AWERB) and were approved by the UK Animals in Science Regulation Unit (ASRU) under the Animals (Scientific Procedures) Act 1986 and authorized by Project License PP6161871 in strict accordance with the Home Office Code of Practice. Experiments were conducted following an authorized protocol reviewed by both the AWERB and the Bioresearch and Veterinary Services (BVS) department at the University of Edinburgh.

Mice used for whole cell recordings and V1 optogenetic stimulation experiments were C57BL/6J females aged between 5 and 8 weeks old at the start of procedures and were obtained from Charles River Laboratories. Mice used for optogenetic silencing were homozygous PV-Cre mice, both males (13) and females (15) aged between 6 and 16 weeks at the start of procedures and were bred and maintained at the University of Edinburgh - founders were originally obtained from The Jackson Laboratory (Strain #:008069). Mice were all co-housed in standard IVC caging with ad libitum water and chow.

### Virtual reality system

For whole cell recordings, mice were head-fixed in a custom toroidal virtual reality system based on previous designs (Dombeck et al., 2010). Mice were free to run on a wheel adapted from the KineMouse wheel design (Warren et al., 2021). The 50 x 80 cm toroidal screen covered a visual field of approximately 280 degrees horizontally and 100 degrees vertically. The virtual tunnel had walls consisting of continuous vertical square gratings. For optogenetics experiments, a V-shaped virtual reality was used consisting of two monitors at a 45-degree angle. The virtual reality environment was generated and controlled by MATLAB-based ViRMEn software (Aronov and Tank, 2014).

### Surgery

#### Head-plate implantation

Aseptic surgical practices were adhered to. Mice were anesthetized with isoflurane (5% for induction, 1.5% to 3% for maintenance), and provided with both local (Bupivacaine, up to 1 mg/kg, s.c.) and general (Meloxicam, 5 mg/kg, s.c.) analgesics at the onset of the surgery. A section of skin and periosteal tissue was removed overlying the dorsal skull, which was then roughened and coated in tissue adhesive (Histoacryl, B Braun, UK). A custom titanium head-plate was adhered to the skull with a combination of tissue adhesive and black dental cement (Paladur, Kulzer, Germany).

#### AAV injections and optic fiber implants

In addition to the head-plate implant, for mice undergoing optogenetics and fiber photometry, an AAV injection and fiber optic cannula implantation took place:

##### Visual cortex activation experiments

9 mice were injected via a burr hole in the right hemisphere with 120 to 160 nl of AAV9-hSyn-ChrimsonR-tdTomato AAV (10^13^ GC/ml) at coordinates (from bregma): 3.9 to 4.2 mm posterior, 2.4 to 2.8 mm lateral, 0.6 mm from the brain surface. A 400 µm 0.37 NA fiber optic cannula was implanted at the same location, with its tip at the cortical surface.

##### Visual cortex silencing experiments

18 mice were injected via a burr hole in the right hemisphere only (10 mice) or bilaterally (8 mice) with 120-160 nl of 1:1 mixture of AAV1-CamKIIa-jGCaMP8m-WPRE and AAV5-hSyn-FLEX-ChrimsonR-tdTomato (7×10^12^ and 10^13^ GC/ml respectively) at the same coordinates as for the activation experiments. A 400 µm 0.37 NA fiber optic cannula was implanted at the same location, with its tip at the cortical surface. In 10 further control mice (6 mice implanted in the right hemisphere only, 4 mice bilaterally implanted), the ChrimsonR AAV was omitted.

*AAV sources:*

- AAV-CamKIIa-jGCaMP8m-WPRE was a gift from Loren Looger (Addgene viral prep # 176751-AAV1; http://n2t.net/addgene:176751; RRID:Addgene_176751).
- pAAV-Syn-FLEX-rc[ChrimsonR-tdTomato] was a gift from Edward Boyden (Addgene viral prep # 62723-AAV5; http://n2t.net/addgene:62723; RRID:Addgene_62723) (Klapoetke et al., 2014)
- pAAV-Syn-ChrimsonR-tdT was a gift from Edward Boyden (Addgene viral prep # 59171-AAV9; http://n2t.net/addgene:59171; RRID:Addgene_59171) (Klapoetke et al., 2014)

#### Recovery

All mice undergoing surgery were allowed to recover in their home cage with mashed food and meloxicam (5 mg/kg) provided for at least 48 hours post-surgery. For mice undergoing AAV injections, recordings would not take place until at least 3 weeks later.

### Mouse habituation

Prior to all recording experiments, mice were trained in daily 20-minute to 1-hour sessions for 4 to 12 days, until they displayed regular locomotion while head fixed. During these training sessions, mice undergoing whole cell recordings were presented with both locomotion-coupled visual flow feedback and the 800 ms fixed-speed visual flow stimuli, so that both types of stimuli were habituated to (and therefore not novel) prior to the recordings. Mice undergoing optical experiments were presented only with visuomotor coupling during habituation.

### Craniotomy and whole cell recordings

Whole cell recordings were performed as described previously (Jordan, 2021; Jordan and Keller, 2020). Mice were provided with Meloxicam (5 mg/kg, s.c.) and a small 1 mm craniotomy was performed over the right V1 (2.5 mm lateral to lambda) under isoflurane anesthesia. The recording chamber was filled with bath recording solution (126 mM NaCl, 5 mM KCl, 10 mM HEPES, 2 mM MgSO_4_, 2 mM CaCl_2_, 12 mM glucose, brought to pH 7.4 using NaOH; 285 mOsm). Mice recovered from anesthesia for at least 20 minutes prior to recordings, which were performed after regular locomotion resumed.

Micropipettes (5-8 MΩ resistance) were fabricated from filamented borosilicate glass (BF150-86-10, Sutter, California, USA) and filled with intracellular recording solution (130 mM KMeSO_3_, 7 mM KCl, 0.1 mM EGTA, 10 mM HEPES, 4 mM Mg-ATP, 0.5 mM Na_2_-GTP, 4 mM Na_2_-phosphocreatine, 1 mM MgCl_2_, 0.02 mM CaCl_2_, brought to pH 7.3 with KOH; 288 mOsm). All recordings took place in current clamp. Data were Bessel low-pass filtered at 10 kHz using a MultiClamp 700B amplifier (Molecular Devices, California USA) and digitized at 20 kHz via a National Instruments PCIe-6363 DAQ.

Voltages have not been corrected for the junction potential. Recordings took place between an estimated 80 to 400 µm from the cortical surface. Electrophysiological properties were determined in the first 120 s using current steps ranging from -0.4 nA to 0.3 nA. Neurons with a spike half-width < 0.7 ms or an input resistance > 150 MΩ were excluded - potential indicators of interneurons (Gentet et al., 2010). In total, 57 neurons were recorded across 32 mice, and three neurons were excluded. Series resistances ranged between 10 and 110 MΩ (mean ± SD = 69 ± 20 MΩ) at the start of recording. For all data included, average V_m_ during stationary periods (and no current injection) was more hyperpolarized than -50 mV.

#### Sequence of stimuli during whole cell recordings

At the start of each recording, 800 ms fixed-speed visual flow stimuli were presented as motion in the virtual tunnel walls, in either the temporonasal or nasotemporal direction in a pseudorandom sequence with a 6.25 ± 0.9 s intertrial interval, during which the gratings remained static, and locomotion did not drive feedback; this period lasted up to 5 minutes. Next, a 5-minute period of visuomotor coupling was presented in which wheel rotation was coupled to linear displacement in the virtual tunnel, generating locomotion-driven visual flow feedback (note: for locomotion-driven recordings, the walls remained static gratings regardless of locomotion during this period). Finally, the fixed-speed visual flow stimuli were presented again for up to 5 minutes. Mice were exposed to both conditions (locomotion-coupled visual flow and fixed-speed stimuli) before and between recordings to limit the impact of novelty.

#### Locomotion-coupled current injection

During visuomotor coupling, locomotion speed was used to control the injected current. In feedback- or locomotion-driven neurons, the gain between locomotion speed and depolarizing current injection was positive to drive firing during feedback, and usually zero (but in some cases, mildly hyperpolarizing) during stationary periods to prevent spiking. In stationary-driven neurons, the gain was negative: hyperpolarizing current scaling with locomotion prevented spiking, while a tonic depolarizing current during stationary periods drove firing. In both cases, a maximum current was set to prevent excessive firing rates, and the gain and maximum were manually adjusted online to ensure reasonable spiking and generate variance across neurons. Neurons were randomly assigned to feedback-driven or stationary-driven before recordings began. If more than one recording was obtained from a mouse, the type of current injection would switch between successive recordings.

### Fiber photometry and optogenetics experiments

Fiber photometry recordings and concurrent optogenetic manipulations were performed with a Neurophotometrics FP3002 system (Mbf Bioscience) using Bonsai software. Pulses of 50-60 µW 470 nm and 410 nm excitation light were interleaved, and green and red fluorescence were recorded simultaneously, with 410 nm (isosbestic) and 470 (calcium fluorescence) channels each recorded at 30 Hz. A branching 200 µm core, 0.37 NA fiber optic patch cord (NPM-BPC-4, Mbf Bioscience) was used to both record fluorescence and deliver optogenetic excitation light. Photometry and behavioral data were aligned using a TTL pulse recorded alongside the behavioral data, which evoked infrequent activation of a red LED recorded using a separate branch of the patch cord.

The following parameters for optogenetic stimulation were used:

#### In V1 silencing experiments

the 635 nm FP3002 optogenetic laser was pulsed at 10 Hz with a 5 to 8 ms pulse width, and a power of 3 to 9 mW at the end of the fiber patch cord. This stimulation occurred either in isolation during visuomotor coupling experience (every 30 s) to test the effect on visual cortex calcium activity (separate recording sessions, with a stimulation duration of 1 s) or beginning 0.5 s prior to the onset of visuomotor mismatch stimuli (which occurred every 10.5 ± 3.5 s), on 25% of trials (total stimulation duration of 1.5 s). In some recordings where calcium activity was not concurrently recorded, a 625 nm fiber-coupled LED (M625F2, Thorlabs) was used to excite ChrimsonR, with the same parameters as above.

#### In V1 activation experiments

The 635 nm FP3002 optogenetic laser was pulsed at either 5 Hz, 10 Hz, or 20 Hz, with a 10 ms pulse width, for 1 s, and a power of 5 to 7 mW at the end of the fiber patch cord. This occurred with an interval of 30 s, uncoupled to any other event. During these experiments, mice experienced visuomotor coupling.

### Pupillometry

During all photometry recordings, the right mouse eye was illuminated with an infrared LED array and recorded at 25 Hz on a CMOS camera (DMK 42BUC03, The Imaging Source) via Bonsai.

## Data analysis

All data analysis was performed using custom written code in MATLAB R2023a.

## WHOLE CELL RECORDING DATA ANALYSIS

### Quantification of spiking during visuomotor coupling

The membrane potential (V_m_) trace was median filtered with a 40 ms window to remove fast spike transients and isolate the subthreshold V_m_, and this filtered trace was subtracted from the original V_m_ trace to isolate spikes, detected as peaks with prominence > 15 mV. Locomotion speed during each spike was recorded, with feedback spikes defined where speed > 4 cm/s and stationary spikes where speed < 1 cm/s. Average rates were obtained by dividing the number of spikes meeting each condition by the number of seconds locomotion also met it. The locomotion speed versus firing rate relationship (**Figure 1E**) was computed similarly, using a series of locomotion speed thresholds. To compare evoked firing rates to physiological rates, the peri-stimulus spike histogram was computed for fixed-speed visual flow stimuli presented during locomotion before visuomotor coupling (**Figure 1F**, gray data), and the maximum firing rate (across 0.25 s bins within the stimulation period) was quantified for each neuron.

### Locomotion-V_m_ correlations

Cross-correlations were calculated between subthreshold V_m_ and locomotion speed for the periods before and after visuomotor coupling. Stationary points (locomotion speed < 1 cm/s) were excluded so that large V_m_ fluctuations of the quiescent state did not influence the correlation. Visual flow stimulus periods, and 1 s following, were also excluded to prevent onset/offset responses affecting the cross-correlation. To assess changes in locomotion-V_m_ correlation (e.g., **Figures 2 and 3**), the cross-correlation was averaged across time lags of -1 to 2 s (positive lags where locomotion speed leads V_m_).

### Responses to visual flow stimuli

The V_m_ response to each stimulus was first baseline subtracted using the average V_m_ in the 1 s immediately preceding the stimulus. Responses were averaged across trials where the average locomotion speed exceeded 4 cm/s during visual flow if the neuron had at least 3 such trials. The response was averaged in the 1 s window starting 125 ms after stimulus onset. Vis_high_ neurons were determined where this average response exceeded 2 mV, and the remainder of neurons were considered Vis_low_ neurons.

### Changes in visual flow response

Multiple measures of change in visual flow response were analyzed (**Figure S3B**). These were calculated from different subsets of visual flow stimuli dependent on the direction of visual flow (nasotemporal or temporonasal), and the behavior of the animal (locomoting: > 4 cm/s average speed during visual flow; stationary: < 0.5 cm/s average speed during visual flow). Response changes were calculated as the difference in average response across trials before vs after visuomotor coupling.

Since changes in baseline V_m_ could account for some variance (∼20%) in these changes, for one measure of response change analyzed in **Figure S3B**, residuals were taken from the linear fits between baseline V_m_ change and changes in response to nasotemporal visual flow during locomotion. Fits and residuals were calculated separately for Vis_low_ and Vis_high_ neurons, and in the latter case, the visual flow response change was first converted to a percentage change owing to the wider variation in starting response values.

The direction selectivity index (DSI) for each neuron was calculated before and after visuomotor coupling:

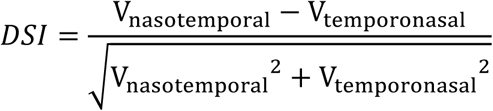

Where V_nasotemporal_ and V_temporonasal_ are the average V_m_ responses to nasotemporal and temporonasal visual flow during locomotion respectively.

### Visuomotor slope analysis

For visuomotor relationships (**Figure 3B-C**), locomotion-V_m_ correlation values were plotted against the average response to nasotemporal visual flow stimuli during locomotion. For each dataset, the two responses were first z-scored, and the slope of the linear fit between them was taken as the visuomotor slope.

### Calculation of electrophysiological variables

#### Change in spike amplitude

Spike amplitudes were used as a correlate of changes in recording quality imposed by changes in series resistance: spike amplitudes were measured in the first visual flow stimulus period (plus 30 s into the visuomotor coupling period) and the second (plus the final 30 s of coupling). Spike amplitudes reduced by 7 ± 8 mV on average across the recording (13 %), with no difference between stationary- and feedback-driven neurons (p = 0.83, unpaired t-test).

#### Resting

*V_m_:* V_m_ was averaged across all datapoints during which the locomotion speed was below 1 cm/s and visual flow was not presented, in the period before visuomotor coupling and the period after. The difference between these two values was taken as the change in resting V_m_.

#### Input resistance

Measured from current steps applied at the beginning of the recording. The total resistance was calculated as the slope of the linear fit between current step size and the average V_m_ response 25 to 125 ms after current step onset. Series resistance was estimated as the slope of the linear fit between current step size and the voltage response in the first 1 ms. Input resistance was calculated as the difference between total resistance and series resistance.

#### Spike threshold

For each spike exceeding 20 mV amplitude evoked during zero current injection, the V_m_ was quantified at the peak of the second derivative of the spike waveform, and the 10^th^ percentile of all values was taken as the spike threshold for the neuron.

#### Spike half-width

For each spike in the initial current-step series exceeding 20 mV in amplitude, threshold was taken as the V_m_ at the peak of the second derivative of the waveform (threshold), and amplitude as the difference between the peak V_m_ and the threshold. Half-width was the duration that the waveform exceeded half this amplitude, and the median was taken across all spikes.

##### Locomotion-related depolarization (‘Depolarization_loco_’)

the V_m_ trace was averaged over all pre-coupling datapoints with locomotion speed < 1 cm/s and no visual flow presented to calculate the stationary V_m_, and over datapoints with locomotion speed > 4 cm/s and no visual flow presented to calculate the locomotion V_m_. Locomotion*-*related depolarization was calculated by subtracting the stationary V_m_ from the locomotion V_m_.

### Multiple linear regression analyses

To determine how much variance was explained in locomotion-V_m_ correlation changes by various predictors (e.g., **Figure 2B and Figure S1B-C**), multiple linear regression was used. Predictor variables and response variables were z-scored. Multiple regression was performed including one to N of the variables in each instance, in all possible combinations. The variance explained by a given variable was calculated as the average increase in R^2^ for models including that variable relative to models with the same combination of predictors but in absence of that variable. Weights for a given variable were computed by averaging coefficients across all models including that variable. The significance of each predictor was determined using stepwise linear regression with all predictors included.

### Variance explained in visual response changes by spiking (Figure S3)

Linear models were fitted between four different measures of neuronal activity during visuomotor coupling (feedback firing rate, feedback spike count, stationary firing rate, and stationary spike count), and 6 different measures of locomotion-related or visual flow response change (see section on ‘visual flow response changes’). For each combination of response changes and neuronal activity measures, a linear model was fitted, with different weights fitted for Vis_low_ and Vis_high_ neurons.

## DESCRIPTION OF THE TWO-PHOTON IMAGING DATASETS

The two-photon imaging datasets were described in detail previously (Jordan and Keller, 2023), but in brief, three datasets were analyzed:

**Figures 4-5 and S4-6**: jGCaMP8m was imaged in layer 2/3 neurons of V1 in NET-Cre mice. In some mice, ChrimsonR was also expressed in LC axons. During imaging sessions, first, 1 s duration fixed-speed visual flow stimuli were presented as motion in the virtual tunnel walls. Between these stimuli, the walls would remain as static gratings, regardless of locomotion. The direction of visual flow stimuli was either temporonasal or nasotemporal, presented in a pseudorandom sequence. During the subsequent 5–10-minute visuomotor coupling period, locomotion generated visual flow feedback. During this time, an optogenetic laser was used periodically to stimulate LC axons in the imaging window (1 s bouts of 20 Hz). Finally, the fixed-speed visual flow stimuli were presented again. Imaging sessions were only analyzed if mice were locomoting above a threshold of 4 cm/s at least 20% of the time during visuomotor coupling.

**Figures 6A-C**: GCaMP6s was imaged in LC axons in various cortical locations (sensory cortex and secondary motor cortex) in NET-Cre mice. During imaging sessions, visuomotor feedback was presented interspersed with visuomotor mismatch stimuli (1 s pauses in the tunnel walls).

**Figures 6D-E**: jGCaMP8m was imaged in layer 2/3 neurons of V1 in NET-Cre mice. During imaging sessions, visuomotor feedback was presented interspersed with visuomotor mismatch stimuli. In some mice, ChrimsonR was also expressed in axons of the LC. During a subset of mismatch events, LC axons were stimulated optogenetically throughout the stimulus. These mismatch stimuli have been excluded from analyses.

## ANALYSES OF VISUOMOTOR PLASTICITY IN CALCIUM ACTIVITY

### Quantification of feedback activity level

Feedback activity level was quantified in the visuomotor coupling period as the difference in average ΔF/F between all points where locomotion speed exceeded 4 cm/s, and all points where locomotion speed was below 0.5 cm/s. Feedback-active neurons were defined as neurons in the top 20% of feedback activity levels, and stationary-active neurons were defined as neurons in the bottom 20% (**Figure 4C**).

### Calcium responses to visual flow stimuli

The response to each nasotemporal stimulus was baseline-subtracted by the average ΔF/F in the 1 s preceding it. Responses were then averaged across trials from the periods before or after visuomotor coupling, where the average locomotion speed during the stimulus exceeded 4 cm/s, if at least 3 such trials existed. The average response was computed in the 2 s window starting 0.33 s after stimulus onset. Responses were corrected by subtracting average responses to sham onsets (evenly spaced every 6 s throughout the recording), selected by average locomotion speed in the same manner.

To define neurons by visual responsivity, responses were averaged across time periods before *and* after the visuomotor coupling (**Figure 4D-F**).

### Calcium responses to feedback onsets and locomotion onsets

Locomotion onsets were identified where locomotion speed crossed a threshold of 2 cm/s, selecting those where average speed was below 0.5 cm/s in the 2 s beforehand and exceeded 2 cm/s in the 1 s after. These were termed feedback onsets during visuomotor coupling (where locomotion evoked visual flow feedback) and locomotion onsets in the periods before and after. Responses were baseline-subtracted using the average ΔF/F in the -2 to -1 s window before onset, then averaged across trials (only where at least 3 valid trials existed). For each neuron, the average response was computed over the 5 s window starting 0.33 s after locomotion onset.

To define neurons by locomotion responsivity, responses were averaged across time periods before *and* after the visuomotor coupling (**Figure S4**).

To calculate changes in feedback onset response, average responses were computed in the first 2.5 minutes of the visuomotor coupling period (‘early’) and the last 2.5 minutes (‘late’). The absolute late response was subtracted from the absolute early response to calculate the absolute feedback onset response changes used in **Figure 5**. In **Figure 5C**, response changes were converted to a change index for each neuron:

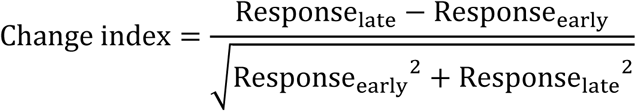

### Calcium responses to optogenetic laser stimulation (Figure 5D-E)

For each neuron, calcium responses to each bout of optogenetic laser stimulation were first baseline subtracted using the average ΔF/F in the 1 s before stimulation onset, then averaged across trials. Only trials in the first 2.5 minutes of visuomotor coupling, and trials during feedback (in which average locomotion speed during the stimulation exceeded 4 cm/s) were included. Each neuron’s average response was then corrected by subtracting an average response to sham onsets (at periodic 6 s intervals) selected in the same way, and the mean response to optogenetic stimulation was calculated over a 2 to 8 s window from onset.

### Relationship between feedback activity level and change in feedback onset response

Each dataset (LC-stimulated and control) was split into deciles grouped by feedback activity level. For each subset, the change in absolute feedback onset response (early vs late in visuomotor coupling) was averaged and plotted against the median feedback activity level (**Figure 5B**).

When comparing this relationship between neurons with high and low responses to LC stimulation (**Figure 5F-G**), broader groupings were used to allow further splitting: 9 partially overlapping subsets grouped by feedback activity level, each representing 20% of the data (the first spanning feedback activity level > 0^th^ and < 20^th^ percentile, the second > 10^th^ and < 20^th^ percentile, and so forth). Within each subset, neurons were further split into those with high average responses to LC stimulation (LC stimulation_high_) when they were in the top third, and those with low responses (LC stimulation_low_) when they were in the bottom third. Remaining neurons were classed as ‘medium’ responses (‘Med.’).

### Relationship between feedback activity levels and changes in visuomotor slope

Each dataset (LC-stimulated and control) was split into quintiles grouped by feedback activity level. For each subset, the slope of the linear fit between visual flow responses and locomotion onset responses was computed using robust fitting, before and after visuomotor coupling. The change in slope (after-before) was plotted against the median feedback activity level of the subset (**Figure 4H-I**).

### Relationship between visual flow responsivity and changes in locomotion onset response

Each dataset (LC-stimulated and control) was split into 5 quintiles grouped by visual flow response. For each subset, the correlation between feedback activity levels and changes in locomotion onset response (before vs after coupling) was computed using robust fitting (**Figures 4E and S5D**), and these coefficients were plotted against the subset’s median visual flow response (**Figures 4F and S5E**).

A similar analysis assessed the relationship between locomotion responsivity and changes in visual flow response, with quintiles instead grouped by locomotion onset responses and correlations computed for each subset between feedback activity levels and changes in visual flow response (**Figure S4**).

## PUPILLOMETRY ANALYSES

### Extraction of pupil diameter from eye-tracking videos

All pupillometry recordings were from the right eye, ipsilateral to the cortical imaging/stimulation sites. The eye-tracking video was manually cropped around the pupil, and a threshold was set by manually tracing the pupil border in three frames of the series and averaging across the 2^nd^ percentile of the surrounding (lightest) pixels and the 98^th^ percentile of the pupil (darkest) pixels. This threshold was applied to binarize all images; the pupil was then detected as an ellipsoid fit to the largest connected component, with its longest axis taken as the diameter.

### Pupil data pre-processing

Blinks/artifacts were detected where the time differential of the pupil diameter exceeded a threshold of 1 SD. Values from 1 sample before to two samples after each blink were removed and the trace was then z-scored using the mean and standard deviation of the pupil diameter during stationary behavior (< 1 cm/s locomotion speed). Mismatch events with missing values in the 1 s preceding to 2.5 s after mismatch onset were excluded. The time derivative of the pupil diameter was analyzed by first smoothing the pupil diameter in a 2-sample window, then differentiating the trace before mean-smoothing within a 0.33 s window. Outliers in the derivative exceeding a threshold of 0.1 z were removed, before z-scoring the trace. In all datasets, recordings with more than 10% of all diameter/derivative values missing were deemed low quality and not analyzed further.

### Relationship between calcium activity and pupil dilation responses to mismatch (Figures 6A-E)

To assess the trial-by-trial relationship between V1 neuronal calcium and pupil diameter responses to mismatches, mismatch trials with average locomotion speed exceeding 1 cm/s during the stimulus were identified. Both calcium responses (per neuron) and pupil diameter responses were baseline-subtracted using the average over the 1 s before the stimulus. Per trial, calcium responses were averaged over the 1 s window starting 0.33 s after mismatch onset and then across all neurons, while the slower pupil responses were averaged over the 1.5 s window starting 1 s after onset. Correlation coefficients between the pupil and averaged calcium responses were then calculated across trials for each site with at least 9 valid mismatch trials.

For LC axon analyses, processing differed in several respects. Active axonal regions of interest (ROIs) were identified as those with a signal-to-noise ratio exceeding 2 (computed as the average FFT power between 0.05 and 1 Hz relative to that between 3 and 8 Hz), and their ΔF/F traces were smoothed within a 150 ms window before averaging across ROIs, to remove high-frequency noise. LC axonal and pupil derivative responses to mismatch were both analyzed in the window 0.66 to 1.66 s after mismatch onset. As well as correlating LC axonal and pupillometric responses across trials within an imaging session, per-trial responses were concatenated from across sessions and divided into quintiles by the size of the pupil derivative response; pupil diameter and LC axon responses were then averaged across concatenated trials for each quintile to generate the plots in **Figure 6B**.

### Response onset delays

For onset-delay analysis between V1 layer 2/3 calcium and pupillometric mismatch responses, only trials with locomotion speed exceeding 4 cm/s during the mismatch event were used. For each session with at least 3 such trials, the calcium response was averaged across trials per neuron and then across neurons exceeding a 1% ΔF/F average mismatch response within each imaging site. Per site, the average V1 calcium and pupil-diameter responses were each divided by the standard deviation over the 2 s before mismatch onset, and the response onset was taken as the first point exceeding +2 standard deviations after onset, where it remained above this threshold for at least the next 0.5 s.

### Pre-processing of fiber photometry data

First, decay of the photometry signals (caused by photobleaching) was removed from both the 410 nm channel (i.e., the isosbestic) and 470 nm (calcium fluorescence) channels by fitting a third order exponential to each and subtracting the fit. The isosbestic channel was then fit to the 470 nm channel using a linear model, and the fit was subtracted from the 470 nm trace to reduce components of the signal caused by hemodynamics and movement artifacts. The resulting trace was then z-scored.

### Analyses of optogenetic activation of PV interneurons (cortical silencing)

Some mice had bilateral implants (8 PV-stimulated mice, 4 controls), while others had unilateral implants in the right V1 (10 PV-stimulated mice, 6 controls). Given no discernable difference was found for bilateral and unilateral stimulation on the ipsilateral pupil, we pooled results from both.

#### Cortical response to PV activation (**Figure 6I**)

Calcium activity was analyzed from the cortical site ipsilateral to the recorded pupil. Onsets of optogenetic laser stimulation bouts were identified (presented alone, and when beginning 0.5 s prior to visuomotor mismatch stimuli). The cortical calcium response on each stimulation was baseline subtracted for the 1 s prior to laser onset, before averaging across all stimulations. Responses were then averaged in the 0.5 s window starting 0.33 s after laser onset. ChrimsonR-injected mice with cortical responses to stimulation above -0.2 z were classified as showing failed suppression.

#### Pupil responses to mismatch (**Figure 6J-K and S7D-E**)

Pupil responses to the mismatch stimuli were analyzed similarly to that described above for the 2-photon imaging datasets, except that the baseline subtraction window was 1 s to 0.5 s prior to mismatch stimulus onset (to avoid the period coinciding with optogenetic stimulation), and mismatch trials were selected where locomotion speed during the stimulus exceeded a higher threshold of 10 cm/s, to ensure large pupil responses and reduce variance due to locomotion speed.

### Responses to optogenetic stimulation of V1 neurons

Optogenetic stimulation trials with missing pupil diameter values within 2 s before to 5 s after stimulation, or with locomotion onsets occurring within 3 s before to 3 s after stimulation onset were excluded. Per-trial pupillary responses were baseline subtracted by the average value in the 1 s prior to stimulus onset, and the response was averaged across trials for each stimulation frequency (**Figure 6F**). Mean responses were quantified in a time window from 0.6 to 2.6 s after stimulation (**Figure 6G**). To quantify behavioral effects of stimulation, changes in locomotion speed following stimulation were quantified similarly (**Figure S7C**). The onset of pupil dilation following optogenetic stimulation in **Figure S7B** was quantified using the average response to 10 Hz and 20 Hz stimulation, as described above for ‘response onset delays’.

### Statistics

All comparative statistical tests are detailed in **Statistical table S1** for electrophysiology experiments, and **Statistical table S2** for optical experiments.

#### Whole cell recordings, group average comparisons

Standard deviations were first tested for a significant difference with a Bartlett test. Where they did not differ significantly, means were compared with an unpaired t-test (or paired t-test for paired data); where they did, group medians were compared with a rank-sum test (or signed-rank test for paired data). For comparisons between four groups defined by visual response type (Vis_low_ and Vis_high_) and current injection type (feedback-driven and stationary-driven) (e.g., **Figures 2E and 3A**), a two-way ANOVA was performed first, and testing between individual groups followed only if any p-value was below 0.05.

#### Box plots and violin plots

The box represents the inter-quartile range (IQR) and median, and the whiskers represent the 10^th^ and 90^th^ percentiles. Violin plots were generated using the ‘daviolinplot.m’ function (Karvelis and oyvindlr, 2024), and the distributions are normalized in each case by the maximum value.

#### Testing the significance of differences in correlation between two groups (whole cell recordings) (e.g., **Figures 2F and 3C**)

Neurons would be permuted 10000 times with respect to group identity (Vis_high_/Vis_low_ or stationary-driven/feedback-driven). For each permuted dataset, the absolute difference in correlation coefficient between the two groups was calculated; the actual difference was deemed significant if it exceeded the 95^th^ percentile of this distribution.

#### Testing the significance of linear fits or correlations across subsets of calcium imaging data

To compare relationships across data subsets (e.g., **Figures 4F, 4H-I, 5B-C, 5F, S4C, S5E and S6D**), two tests were used:

1. **Permutation test**: Spearman’s rank correlation coefficient (rho) was computed for the actual data, then neurons were shuffled within each imaging site with respect to the subset-defining variable (e.g., feedback activity level, as in **Figure 4H**) and rho recomputed for the permuted dataset; this was repeated 1000 times. P_shuff_ was the proportion of permuted absolute correlation values exceeding the actual absolute value, with significance at P_shuff_ < 0.05.
2. **Hierarchical bootstrap test**: To account for the data’s nested structure and ensure relationships did not depend on a small number of imaging sites, we performed hierarchical bootstrapping (Saravanan et al., 2020). Control and LC-stimulated datasets would be bootstrapped at the level of imaging sites and neurons 1000 times, with replacement, and Spearman’s correlation coefficient would be computed for the relationship for each bootstrapped dataset. 95% confidence intervals (CI_95_) were derived from this bootstrapped distribution as the 2.5^th^ and 97.5^th^ percentiles. Where > 97.5% of correlation values were above or below zero (P_boot_ < 0.025), significance was determined.

The same approaches were used to test the significance of differences in linear fits or correlations between LC stimulated and control groups (e.g., **Figure 4I, 5B, and S6D**), except that in the permutation test, imaging sites would be shuffled 1000 times with respect to experimental group.

## Supplementary Figures

**Figure S1.**
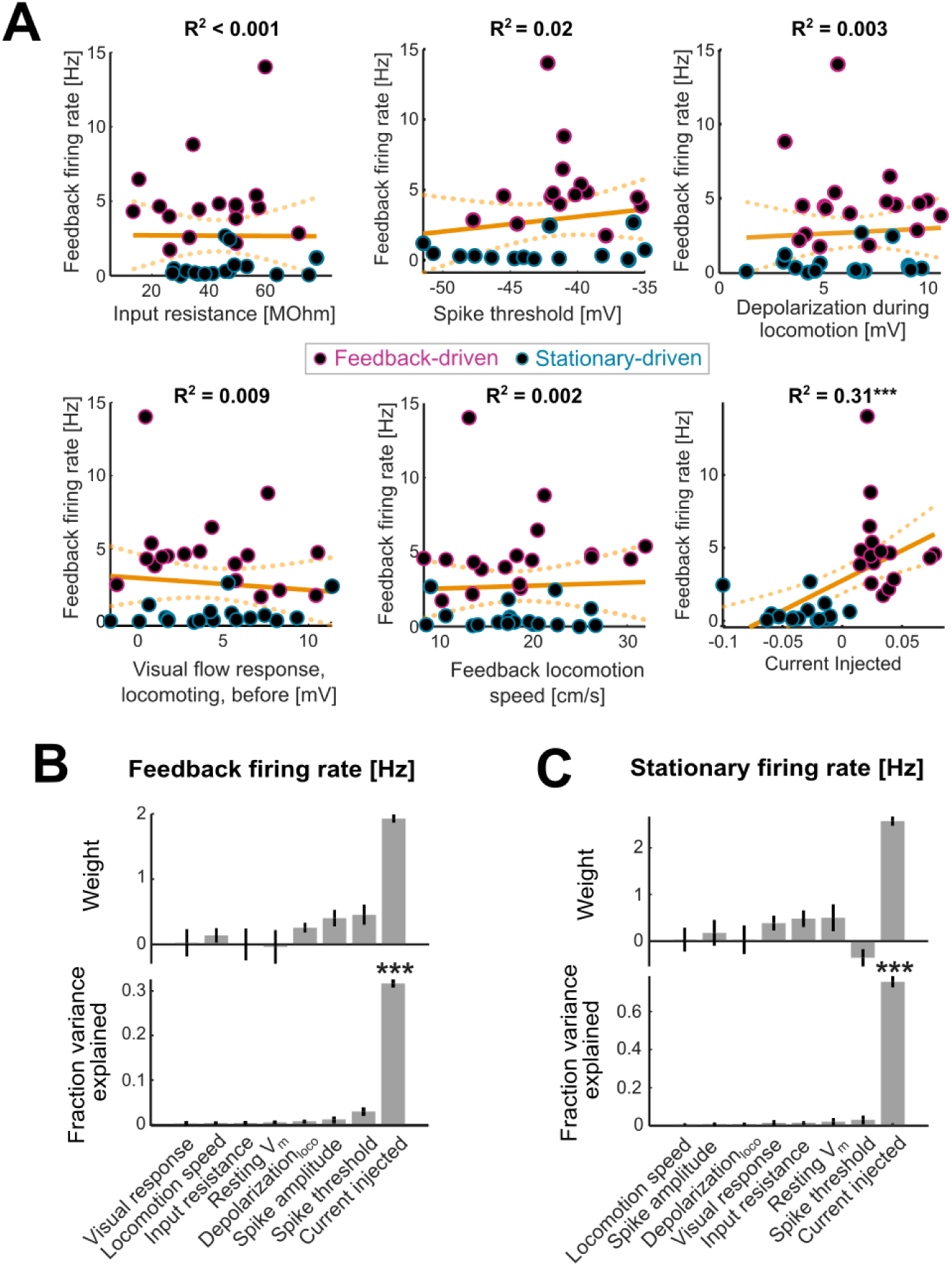
Firing rates during visuomotor coupling are determined primarily by current injection. **(A)** Scatter plots between the firing rate during feedback and various factors that could theoretically impact this firing rate. The ‘current injected’ variable in the last plot refers to the current injected during locomotion at 15 cm/s (a measure of the current injection gain). Orange lines indicate the linear fit to the data, with dotted curves indicating the 95% confidence intervals. **(B)** Average weights and variance explained for eight different predictors of the feedback firing rate across all neurons, computed using multiple linear regression – see Methods. Error bars show standard deviation. Asterisks indicate significance determined by stepwise linear regression. Visual response = average subthreshold response to nasotemporal visual flow stimuli (measured during locomotion, before visuomotor coupling); Locomotion speed = average locomotion speed during visuomotor feedback; Depolarization_loco_ = average depolarization during locomotion relative to stationary periods before visuomotor coupling. **(C)** As for panel B, but for firing rates during stationary behavior during the visuomotor coupling period. The ‘current injected’ variable here refers to the current injected during locomotion at 0 cm/s.

**Figure S2.**
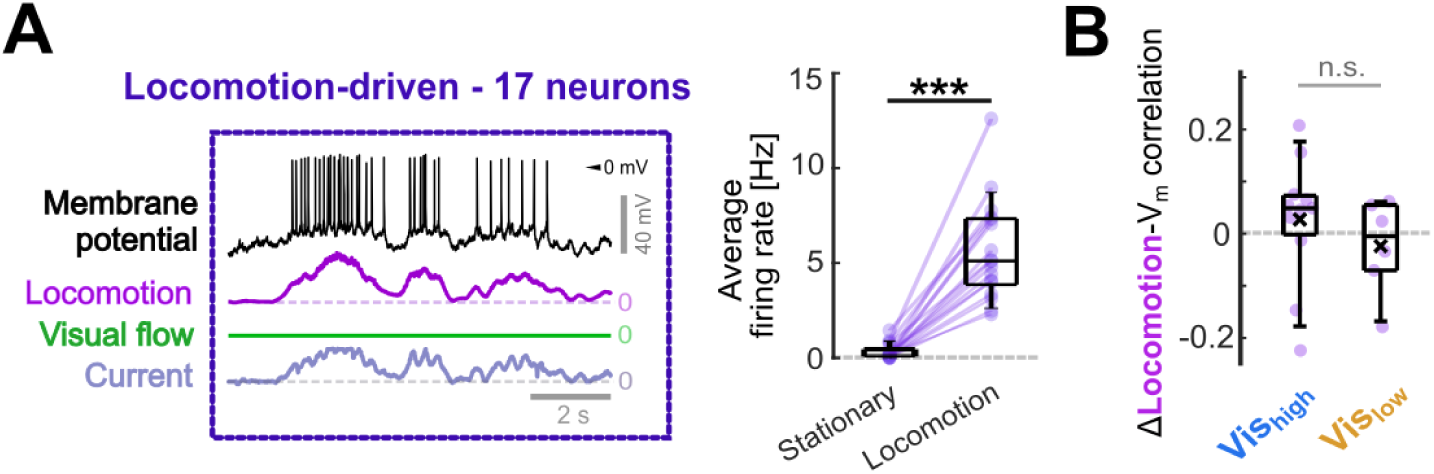
Locomotion-V_m_ changes in neurons where current injection correlated with locomotion without visual flow feedback. **(A)** Example segment of the current injection period for a neuron in a separate group of ‘locomotion-driven’ neurons, where depolarizing current injection scaled with locomotion speed (as for feedback-driven neurons), but visual flow feedback was omitted (walls remained as static gratings). Boxplot shows the average firing rates of these neurons during stationary and locomotion periods. P-value indicates the outcome of paired t-test: p < 0.001; See **Statistical table S1** for details. **(B)** Changes in locomotion-V_m_ correlation (after-before the current injection period) for locomotion-driven neurons. Outcome of comparative test is indicated: p < 0.05; See **Statistical table S1** for details.

**Figure S3.**
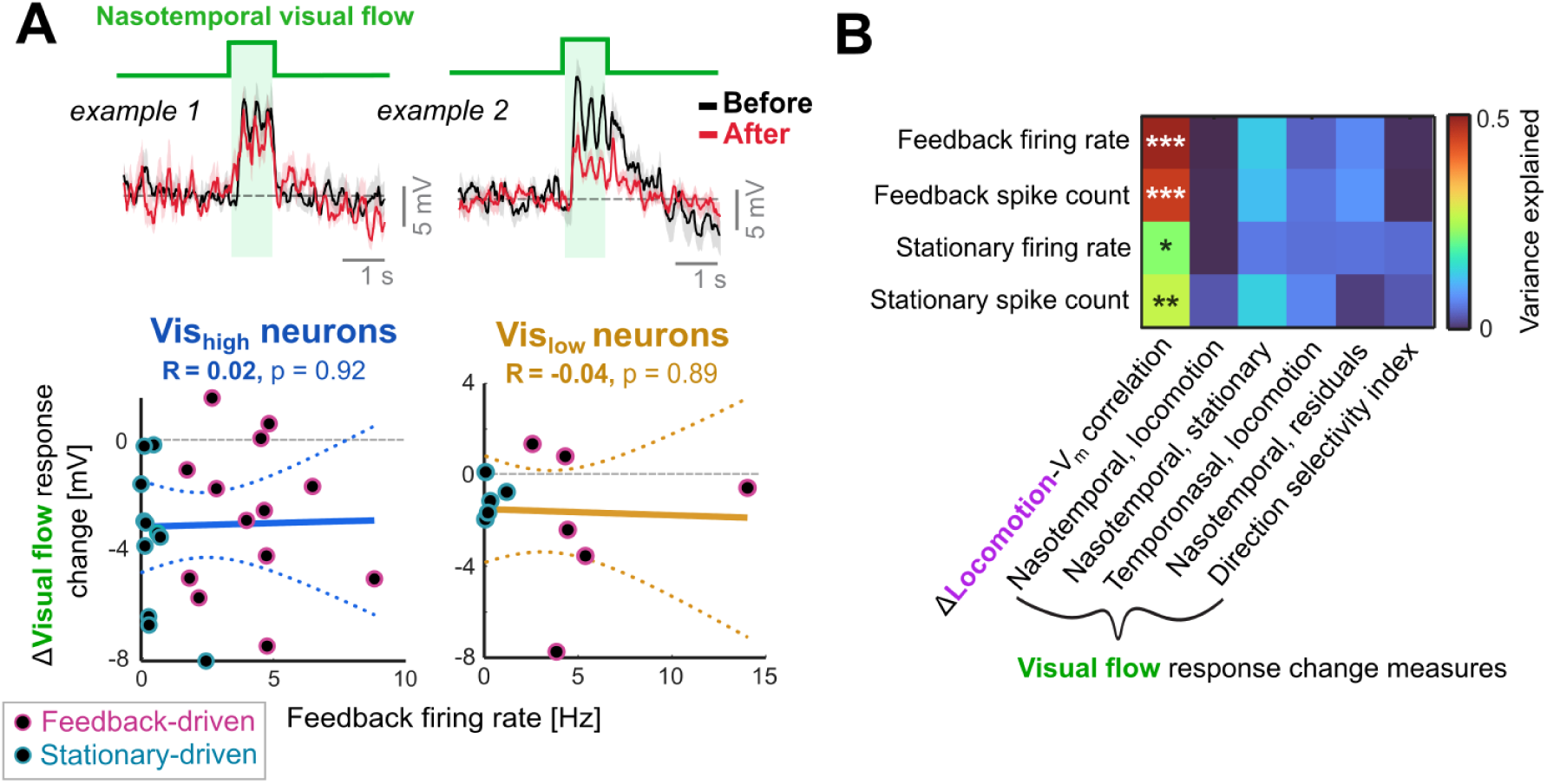
Analyses on changes in visual flow responses in the whole cell recording experiments. **(A)** Top: Average subthreshold responses to nasotemporal visual flow stimuli during locomotion before (black) and after (red) visuomotor coupling for two example Vis_high_ neurons. Shading shows SEM. Bottom: Relationship between feedback firing rates and visual flow response changes (nasotemporal visual flow during locomotion) in Vis_low_ and Vis_high_ neurons. **(B)** Variance explained for various measures of V_m_ response change (x-axis labels) by various measures of neuronal activity during visuomotor coupling (y axis labels). For each pair of predictors and response variables, a linear model was fitted with separate weights for Vis_high_ and Vis_low_ neurons. Asterisks indicate the p-value of the model fit: *: p < 0.05, **: p < 0.01, ***: p < 0.001. ‘Nasotemporal, residuals’ = the residual changes in response to nasotemporal visual flow after fitting these response changes to changes in baseline V_m_, which could account for some variance in response changes - see Methods. Note that significant effects were only found for locomotion-related response changes and were not found for any measure of visual flow response change.

**Figure S4.**
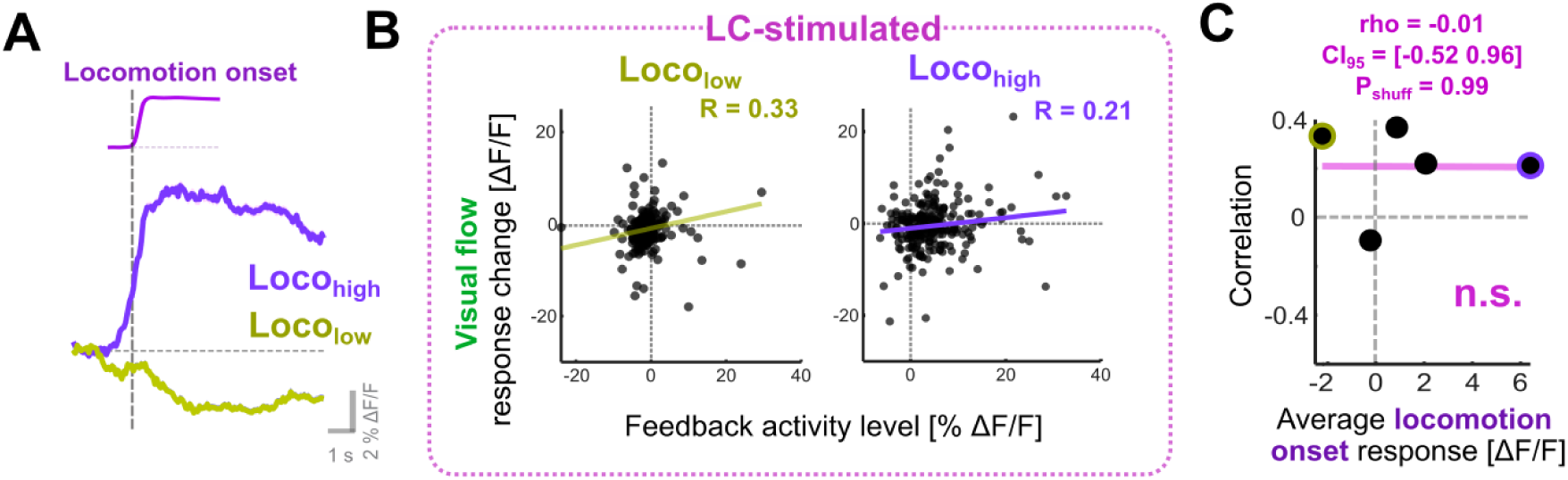
Lack of relationship between locomotion responsivity and visual flow response changes. **(A)** Average calcium responses to locomotion onset were used to identify Loco_high_ and Loco_low_ neurons, which are respectively activated or suppressed by locomotion onset. Traces show average responses across neurons, and shading indicates SEM. **(B)** Correlations between changes in visual flow response (after-before visuomotor coupling), and feedback activity levels in LC-stimulated mice, for neurons defined as Loco_high_ and Loco_low_. **(C)** Correlation coefficients between feedback activity levels and changes in visual flow response (as shown in panel B) plotted against average locomotion onset response for quintiles of the data grouped by locomotion onset responses (see Methods). Note that no subsets show a substantial negative relationship, as would be expected if spiking drove increases in visually driven inhibition.

**Figure S5.**
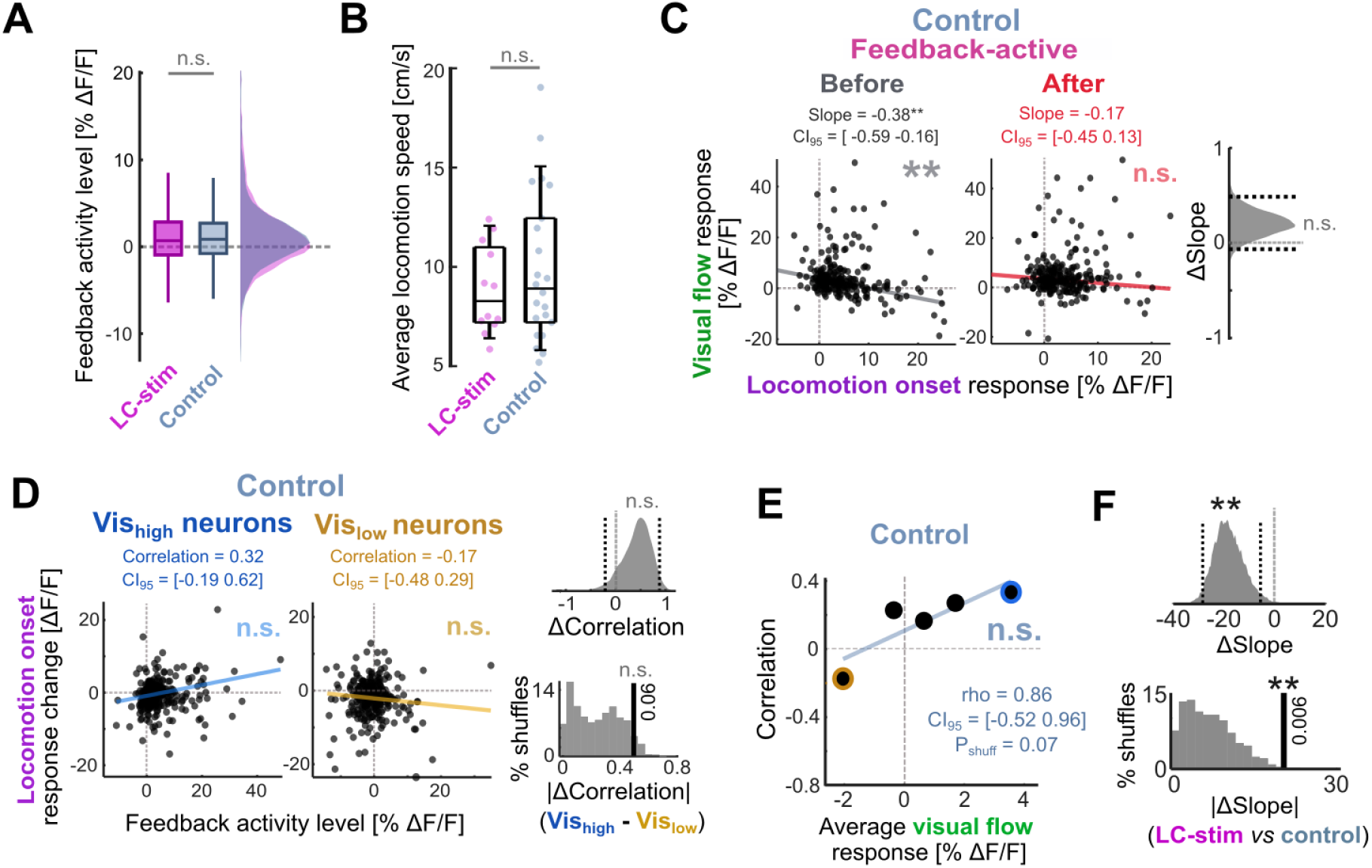
Comparisons of activity-dependent calcium response changes between control and LC-stimulated datasets. **(A)** Comparison of feedback activity levels across LC-stimulated and control datasets. ‘n.s.’ indicates outcome of hierarchical bootstrap test (P_boot_ < 0.05). See **Statistical table S2** for details. **(B)** Comparison of average locomotion speed during visuomotor coupling for LC-stimulated and control imaging sessions. ‘n.s.’ indicates outcome of rank-sum test (p < 0.05). See **Statistical table S2** for details. **(C)** Relationship between visual flow and locomotion onset responses (Z-scored) before and after visuomotor coupling for feedback-active neurons in control mice. Linear fit slopes (visuomotor slopes), 95% confidence intervals (CI_95_), and p-values were obtained by hierarchical bootstrapping (see Methods). Violin plot shows the distribution of slope changes across 1000 hierarchically bootstrapped datasets; black dotted lines indicate CI_95_. **(D)** Locomotion onset response changes (after − before visuomotor coupling) versus feedback activity levels for Vis_low_ and Vis_high_ neurons in control mice. Linear fit CI_95_ and p-values were obtained by hierarchical bootstrapping (see Methods). Violin plot shows the distribution of correlation differences between Vis_low_ and Vis_high_ neurons across 1000 hierarchically bootstrapped datasets, with CI_95_ indicated by black dotted lines. Histogram compares absolute slope difference with those from 1000 datasets permuted with respect to visual flow response (see Methods). **(E)** Correlation coefficients between feedback activity level and locomotion onset response changes (as in panel D) plotted against median visual flow responses for visual-response quintiles. P_shuff_ was derived from permutation analysis; CI_95_ values were obtained by hierarchical bootstrapping (see Methods). **(F)** Violin plot shows the hierarchically bootstrapped distribution of slope differences between control and LC-stimulated datasets (black dotted lines indicate CI_95_). Histogram below compares absolute slope difference with those from 1000 datasets permuted with respect to experimental group (see Methods).

**Figure S6.**
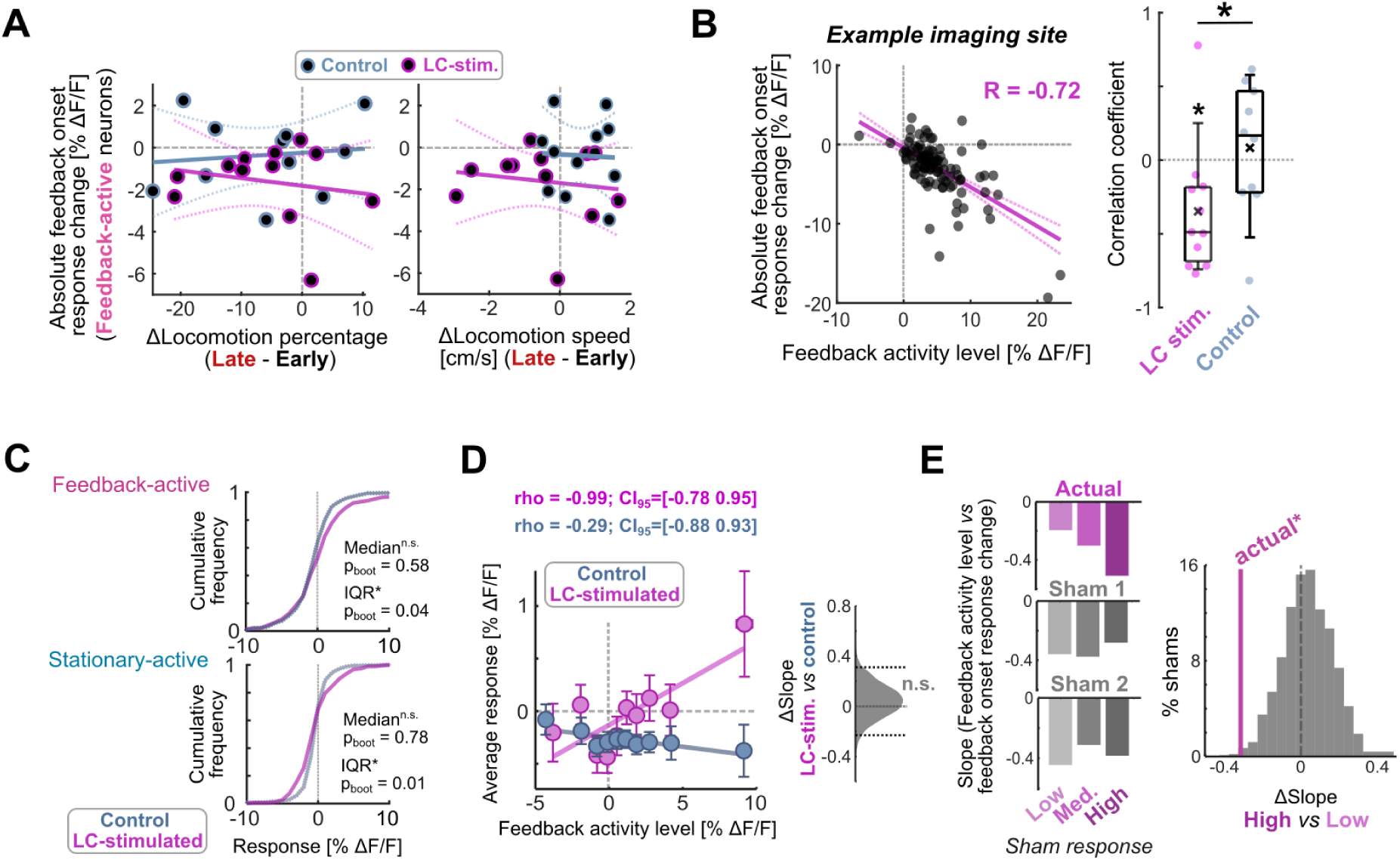
Further analyses on changes in feedback onset response and LC axonal stimulation. **(A)** Average change in absolute feedback onset response (averaged across feedback-active neurons in each imaging site) plotted against the change in locomotion percentage or the change in locomotion speed (late relative to early in the visuomotor coupling period). All correlations were non-significant (p > 0.05). **(B)** Left: Average change in absolute feedback onset response plotted against feedback activity level across neurons for an example imaging site with LC axon stimulation. Right: Correlation coefficients compared across LC-stimulated and control imaging sites for the relationship shown on the left. Asterisks indicate outcome of t-tests: *: p < 0.05. See **Statistical Table S2** for details. **(C)** Cumulative histograms comparing calcium responses to optogenetic laser stimulation in LC-stimulated and control mice for feedback-active and stationary-active neurons. Outcomes of hierarchical bootstrap tests for differences in median and interquartile range (IQR) are shown. **(D)** Mean calcium responses to LC axon stimulation plotted against average feedback activity level (black) for non-overlapping 10% data subsets grouped by feedback activity level. Error bars indicate SEM. Violin plot shows the distribution of slope differences between control and LC-stimulated datasets across 1000 hierarchical bootstraps; dotted lines indicate 95% confidence intervals. **(E)** Linear fit slopes between average feedback activity level and mean changes in feedback onset responses for LC-stimulated data, with neurons grouped by their calcium responses to LC axon stimulation (‘actual’; as in Figure 5F-G), or grouped identically using responses to sham onsets (two examples shown in gray). Histogram shows the differences in slope between neurons with high versus low sham responses (1000 datasets with randomized sham onsets), compared with the corresponding difference based on LC-stimulation responses (actual). Asterisk indicates p < 0.05.

**Figure S7.**
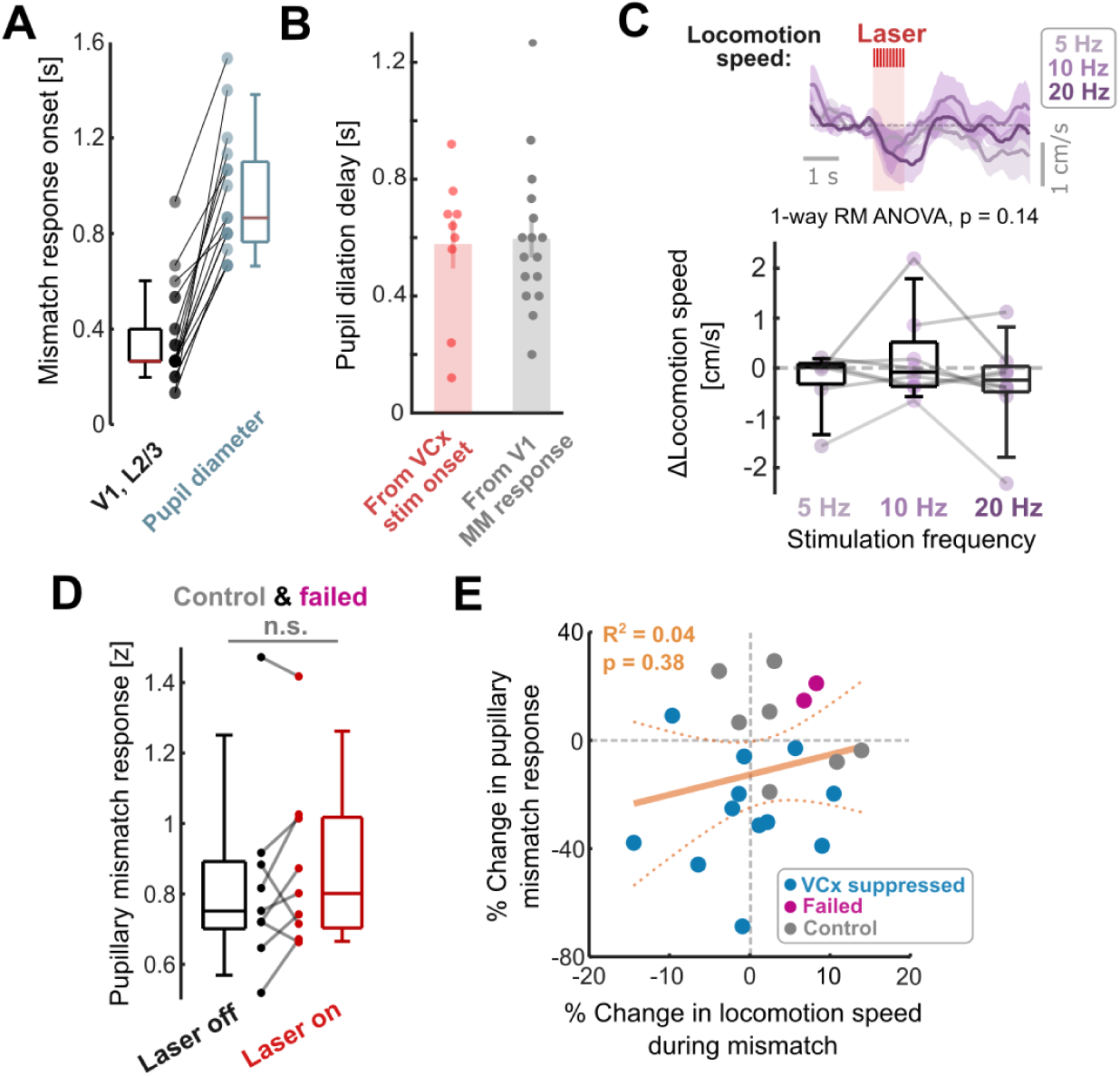
Additional pupillometry analyses. **(A)** Average onset delay for V1 layer 2/3 calcium responses (averaged across mismatch responsive neurons only) and pupillary responses following visuomotor mismatch across imaging sites (see Methods). **(B)** Delay in onset of pupil dilation following optogenetic visual cortex stimulation (VCx stim), and pupil dilation following visuomotor mismatch, relative to the onset of V1 calcium responses recorded concurrently (‘From V1 MM response’), calculated as the difference in values in panel A. Note the similarity in timing. **(C)** Top: Average change in locomotion speed across mice following optogenetic stimulation of visual cortex. Shading shows SEM. Bottom: Comparison of average change in locomotion speed for all mice following visual cortex stimulations at different frequencies. **(D)** Average pupil diameter responses to mismatch between trials with and without the optogenetic laser across all control mice and PV-stimulated mice with failed V1 suppression. P-value indicates outcome of paired t-test (p < 0.05). **(E)** Change in pupillary mismatch response between trials with the optogenetic laser on and off plotted against the change in average locomotion speed during mismatch across the same set of trials.

---

**Statistical table S1.** Whole cell recording analyses.

| Figure panel | Comparison | Mean 1<br>*difference shown for paired* | SD 1<br>(IQR) | Mean 2 | SD 2<br>(IQR) | P value | Test | # Neurons 1 | # Neurons 2 |
| --- | --- | --- | --- | --- | --- | --- | --- | --- | --- |
| 1F | Feedback-driven neurons feedback firing rate vs stationary-driven neurons stationary firing rate [Hz] | 4.75 | 2.79 | 5.27 | 2.36 | 0.28 | Rank sum | 19 | 18 |
| 1F | Feedback-driven neurons stationary firing rate vs stationary-driven neurons feedback firing rate [Hz] | 0.17 | 0.19 | 0.56 | 0.79 | 0.057 | Rank sum | 19 | 18 |
| 1F | Stationary-driven neurons, difference between stationary firing rate and feedback firing rate [Hz] | -4.71 | 1.90 | n/a | n/a | **<10<sup>-8</sup>** | Paired t-test | 18 | n/a |
| 1F | Feedback-driven neurons, difference between stationary firing rate and feedback firing rate [Hz] | 4.58 | 2.74 | n/a | n/a | **<10<sup>-3</sup>** | Signed-rank test | 19 | n/a |
| 2E | Feedback-driven vs stationary-driven neurons, Vis<sub>high</sub>, Locomotion-V<sub>m</sub> correlation change | -0.08 | 0.13 | 0.03 | 0.09 | **0.012** | Unpaired t-test | 13 | 13 |
| 2E | Feedback-driven vs stationary-driven neurons, Vis<sub>low</sub>, Locomotion-V<sub>m</sub> correlation change | 0.13 | 0.13 | -0.09 | 0.08 | **0.012** | Unpaired t-test | 6 | 5 |
| 2E | Vis<sub>high</sub> vs Vis<sub>low</sub>, feedback-driven neurons, Locomotion-V<sub>m</sub> correlation change | -0.08 | 0.13 | 0.13 | 0.13 | **0.003** | Unpaired t-test | 13 | 6 |
| 2E | Vis<sub>high</sub> vs Vis<sub>low</sub>, stationary-driven neurons, Locomotion-V<sub>m</sub> correlation change | 0.03 | 0.09 | -0.09 | 0.08 | **0.021** | Unpaired t-test | 13 | 5 |
| 3A | Feedback-driven vs stationary-driven neurons, Vis<sub>high</sub>, Locomotion-V<sub>m</sub> correlation After | -0.01 | 0.10 | 0.09 | 0.09 | **0.009** | Unpaired t-test | 13 | 13 |
| 3A | Feedback-driven vs stationary-driven neurons, Vis<sub>low</sub>, Locomotion-V<sub>m</sub> correlation After | 0.17 | 0.08 | 0.01 | 0.06 | **0.005** | Unpaired t-test | 6 | 5 |
| 3A | Vis<sub>high</sub> vs Vis<sub>low</sub>, feedback-driven, Locomotion-V<sub>m</sub> correlation After | -0.01 | 0.10 | 0.17 | 0.08 | **0.001** | Unpaired t-test | 13 | 6 |
| 3A | Vis<sub>high</sub> vs Vis<sub>low</sub>, stationary-driven neurons, Locomotion-V<sub>m</sub> correlation After | 0.09 | 0.09 | 0.01 | 0.06 | 0.08 | Unpaired t-test | 13 | 5 |
| S2A | Locomotion-driven neurons, difference between stationary firing rate and locomotion firing rate [Hz] | 5.30 | 2.47 | n/a | n/a | **<10<sup>-6</sup>** | Paired t-test | 17 | n/a |
| S2B | Locomotion-driven neurons, Vis<sub>low</sub> vs Vis<sub>high</sub>, Locomotion-V<sub>m</sub> correlation change | 0.03 | 0.12 | -0.02 | 0.10 | 0.39 | Unpaired t-test | 11 | 6 |

**Statistical table S2.** Optical experiment analyses.

| Figure panel | Comparison | Mean 1<br>difference<br>shown for<br>paired | SD 1 | Mean 2 | SD 2 | P value | Test | # Neurons,<br>sites,<br>animal | # Neurons,<br>sites,<br>animals |
| --- | --- | --- | --- | --- | --- | --- | --- | --- | --- |
| 5E | LC-stimulated vs control, LC-stimulation response [%ΔF/F] (median and IQR shown) | -0.003 | 2.9 | -0.26 | 2.0 | 0.88<br>(Med.)<br>**0.01**<br>(IQR) | Hierarchical bootstrap | 1054 neurons, 11 sites, 5 mice | 2314 neurons, 22 sites, 10 mice |
| 6C | Correlation coefficient between LC axonal and pupil derivative mismatch responses | 0.38 | 0.27 | n/a | n/a | **< 0.001** | Sign-rank test | 34 sites, 12 mice | n/a |
| 6C | Correlation coefficient between LC axonal and pupil diameter mismatch responses | 0.28 | 0.29 | n/a | n/a | **< 0.001** | Paired t-test | 34 sites, 12 mice | n/a |
| 6E | Correlation coefficient between V1 and pupil diameter mismatch responses | 0.14 | 0.22 | n/a | n/a | 0.005 | Paired t-test | 23 sites, 14 mice | n/a |
| 6G | Pupillary response to 5 Hz cortical stimulation [z] | 0.13 | 0.21 | n/a | n/a | 0.17 | Paired t-test | 7 mice | n/a |
| 6G | Pupillary response to 10 Hz cortical stimulation [z] | 0.43 | 0.32 | n/a | n/a | **<0.01** | Paired t-test | 8 mice | n/a |
| 6G | Pupillary response to 20 Hz cortical stimulation [z] | 0.62 | 0.28 | n/a | n/a | **<0.001** | Paired t-test | 8 mice | n/a |
| 6J | Laser on vs laser off, pupillary mismatch response, V1 suppressed mice [z] | -0.15 | 0.16 | n/a | n/a | **0.006** | Paired t-test | 12 mice | n/a |
| 6K | VCx suppressed vs control+failed mice, % change in pupillary mismatch response (laser on vs laser off trials) [%] | 9 | 16 | -26 | 21 | **0.0005** | Unpaired t-test | 12 mice | 9 mice |
| S5A | LC-stimulated vs control, Feedback activity levels [% ΔF/F] | 1.6 | 4.9 | 1.3 | 4.6 | 0.85 | Hierarchical bootstrap | 1196 neurons, 12 sites, 5 mice | 2314 neurons, 22 sites, 10 mice |
| S5B | LC-stimulated vs control, locomotion speed during visuomotor feedback [cm/s] | 8.9 | 2.2 | 9.9 | 3.8 | 0.37 | Rank-sum | 12 sites, 5 mice | 22 sites, 10 mice |
| S6B | Correlation coefficient, feedback activity vs feedback onset response change, LC-stimulated | -0.35 | 0.44 | n/a | n/a | **0.026** | Paired t-test | 11 sites | n/a |
| S6B | Correlation coefficient, feedback activity vs feedback onset response change, control | 0.08 | 0.45 | n/a | n/a | 0.57 | Paired t-test | 10 sites | n/a |
| S6B | Correlation coefficient, feedback activity vs feedback onset response change, LC-stimulated vs control | -0.35 | 0.44 | 0.08 | 0.45 | **0.038** | Unpaired t-test | 11 sites | 10 sites |
| S6C | LC-stimulated vs control, feedback-active, LC-stimulation response [%ΔF/F] (median and IQR shown) | 0.44 | 4.66 | -0.34 | 3.04 | 0.58<br>(Med.)<br>**0.04**<br>(IQR) | Hierarchical bootstrap | 248 neurons, 11 sites, 5 mice | 444 neurons, 22 sites, 10 mice |
| S6C | LC-stimulated vs control, stationary-active, LC-stimulation response [%ΔF/F] (median and IQR shown) | -0.07 | 2.66 | -0.13 | 2.02 | 0.78<br>(Med.)<br>**0.01**<br>(IQR) | Hierarchical bootstrap | 191 neurons, 11 sites, 5 mice | 444 neurons, 22 sites, 10 mice |
| S7D | Laser on vs laser off, pupillary mismatch response, control and failed suppression mice [z] | 0.07 | 0.08 | n/a | n/a | **0.047** | Paired t-test | 8 mice | n/a |

## Notes

### Competing Interest Statement

The authors have declared no competing interest.

### Summary of Updates

Substantial re-write (prior conclusions unchanged); Addition of analyses relating to calcium changes following LC stimulation (Figure 5 and S6C-E); Addition of pupillometry and optogenetics experiments in Figure 6 and S7.

## References

Aronov, D., Tank, D.W., 2014. Engagement of neural circuits underlying 2D spatial navigation in a rodent virtual reality system. Neuron 84, 442–456. 10.1016/j.neuron.2014.08.042

Aston-Jones, G., Cohen, J.D., 2005. AN INTEGRATIVE THEORY OF LOCUS COERULEUS-NOREPINEPHRINE FUNCTION: Adaptive Gain and Optimal Performance. Annual Review of Neuroscience 28, 403–450. 10.1146/annurev.neuro.28.061604.135709

Attinger, A., Wang, B., Keller, G.B., 2017. Visuomotor Coupling Shapes the Functional Development of Mouse Visual Cortex. Cell 169, 1291–1302.e14. 10.1016/j.cell.2017.05.023

Audette, N.J., Schneider, D.M., 2023. Stimulus-Specific Prediction Error Neurons in Mouse Auditory Cortex. J. Neurosci. 43, 7119–7129. 10.1523/JNEUROSCI.0512-23.2023

Bastos, A.M., Usrey, W.M., Adams, R.A., Mangun, G.R., Fries, P., Friston, K.J., 2012. Canonical microcircuits for predictive coding. Neuron 76, 695–711. 10.1016/j.neuron.2012.10.038

Basu, A., Yang, J.-H., Yu, A., Glaeser-Khan, S., Rondeau, J.A., Feng, J., Krystal, J.H., Li, Y., Kaye, A.P., 2024. Frontal Norepinephrine Represents a Threat Prediction Error Under Uncertainty. Biological Psychiatry 0. 10.1016/j.biopsych.2024.01.025

Bear, M.F., Singer, W., 1986. Modulation of visual cortical plasticity by acetylcholine and noradrenaline. Nature 320, 172–176. 10.1038/320172a0

Cole, N., Harvey, M., Myers-Joseph, D., Gilra, A., Khan, A.G., 2024. Prediction-error signals in anterior cingulate cortex drive task-switching. Nat Commun 15, 7088. 10.1038/s41467-024-51368-9

Crapse, T.B., Sommer, M.A., 2008. Corollary discharge across the animal kingdom. Nat. Rev. Neurosci. 9, 587–600. 10.1038/nrn2457

Dombeck, D.A., Harvey, C.D., Tian, L., Looger, L.L., Tank, D.W., 2010. Functional imaging of hippocampal place cells at cellular resolution during virtual navigation. Nat. Neurosci. 13, 1433–1440. 10.1038/nn.2648

Duss, S.N., Wilhelm, M., Marinescu, A.-M., Zhang, R., Helmchen, F., Bohacek, J., Rupprecht, P., 2026. Hippocampal neuronal and astrocytic responses to noradrenaline and natural arousal. eLife 15. 10.7554/eLife.111593.1

Fiser, A., Mahringer, D., Oyibo, H.K., Petersen, A.V., Leinweber, M., Keller, G.B., 2016. Experience-dependent spatial expectations in mouse visual cortex. Nat. Neurosci. 19, 1658–1664. 10.1038/nn.4385

Furutachi, S., Franklin, A.D., Aldea, A.M., Mrsic-Flogel, T.D., Hofer, S.B., 2024. Cooperative thalamocortical circuit mechanism for sensory prediction errors. Nature 633, 398–406. 10.1038/s41586-024-07851-w

Garner, A.R., Keller, G.B., 2022. A cortical circuit for audio-visual predictions. Nat Neurosci 25, 98–105. 10.1038/s41593-021-00974-7

Gentet, L.J., Avermann, M., Matyas, F., Staiger, J.F., Petersen, C.C.H., 2010. Membrane potential dynamics of GABAergic neurons in the barrel cortex of behaving mice. Neuron 65, 422–435. 10.1016/j.neuron.2010.01.006

Gray, S.R., Ye, L., Ye, J.Y., Paukert, M., 2021. Noradrenergic terminal short-term potentiation enables modality-selective integration of sensory input and vigilance state. Science Advances 7, eabk1378. 10.1126/sciadv.abk1378

Hamm, J.P., Shymkiv, Y., Han, S., Yang, W., Yuste, R., 2021. Cortical ensembles selective for context. PNAS 118. 10.1073/pnas.2026179118

Hertäg, L., Sprekeler, H., 2020. Learning prediction error neurons in a canonical interneuron circuit. eLife 9, e57541. 10.7554/eLife.57541

Hippenmeyer, S., Vrieseling, E., Sigrist, M., Portmann, T., Laengle, C., Ladle, D.R., Arber, S., 2005. A developmental switch in the response of DRG neurons to ETS transcription factor signaling. PLoS Biol 3, e159. 10.1371/journal.pbio.0030159

Jordan, R., 2024. The locus coeruleus as a global model failure system. Trends in Neurosciences 47, 92–105. 10.1016/j.tins.2023.11.006

Jordan, R., 2021. Optimized protocol for in vivo whole-cell recordings in head-fixed, awake behaving mice. STAR Protoc 2, 100347. 10.1016/j.xpro.2021.100347

Jordan, R., Keller, G.B., 2023. The locus coeruleus broadcasts prediction errors across the cortex to promote sensorimotor plasticity. eLife 12, RP85111. 10.7554/eLife.85111

Jordan, R., Keller, G.B., 2020. Opposing Influence of Top-down and Bottom-up Input on Excitatory Layer 2/3 Neurons in Mouse Primary Visual Cortex. Neuron 108, 1194–1206.e5. 10.1016/j.neuron.2020.09.024

Karvelis, P., oyvindlr, 2024. povilaskarvelis/DataViz: v3.2.4. 10.5281/zenodo.12749045

Keller, G.B., Bonhoeffer, T., Hübener, M., 2012. Sensorimotor mismatch signals in primary visual cortex of the behaving mouse. Neuron 74, 809–815. 10.1016/j.neuron.2012.03.040

Keller, G.B., Mrsic-Flogel, T.D., 2018. Predictive Processing: A Canonical Cortical Computation. Neuron 100, 424–435. 10.1016/j.neuron.2018.10.003

Klapoetke, N.C., Murata, Y., Kim, S.S., Pulver, S.R., Birdsey-Benson, A., Cho, Y.K., Morimoto, T.K., Chuong, A.S., Carpenter, E.J., Tian, Z., Wang, J., Xie, Y., Yan, Z., Zhang, Y., Chow, B.Y., Surek, B., Melkonian, M., Jayaraman, V., Constantine-Paton, M., Wong, G.K.-S., Boyden, E.S., 2014. Independent optical excitation of distinct neural populations. Nat Methods 11, 338–346. 10.1038/nmeth.2836

Leinweber, M., Ward, D.R., Sobczak, J.M., Attinger, A., Keller, G.B., 2017. A Sensorimotor Circuit in Mouse Cortex for Visual Flow Predictions. Neuron 95, 1420–1432.e5. 10.1016/j.neuron.2017.08.036

Martins, A.R.O., Froemke, R.C., 2015. Coordinated forms of noradrenergic plasticity in the locus coeruleus and primary auditory cortex. Nat Neurosci 18, 1483–1492. 10.1038/nn.4090

Marzo, A., Bai, J., Otani, S., 2009. Neuroplasticity Regulation by Noradrenaline in Mammalian Brain. Curr Neuropharmacol 7, 286–295. 10.2174/157015909790031193

Matarasso, A., Reyes, I.R., Seaholm, E., Cheeyandira, A., Piantadosi, S.C., Li, L., Li, Y., Weinshenker, D., Bruchas, M.R., 2025. Stimulus-Dependent Dopamine Dynamics from Locus Coeruleus Axons. bioRxiv 2025.09.15.676390. 10.1101/2025.09.15.676390

Mikulasch, F.A., Rudelt, L., Priesemann, V., 2022. Visuomotor Mismatch Responses as a Hallmark of Explaining Away in Causal Inference. Neural Comput 35, 27–37. 10.1162/neco_a_01546

Mikulasch, F.A., Rudelt, L., Wibral, M., Priesemann, V., 2023. Where is the error? Hierarchical predictive coding through dendritic error computation. Trends in Neurosciences 46, 45–59. 10.1016/j.tins.2022.09.007

Nestvogel, D.B., McCormick, D.A., 2022. Visual thalamocortical mechanisms of waking state-dependent activity and alpha oscillations. Neuron 110, 120–138.e4. 10.1016/j.neuron.2021.10.005

Neyhart, E., Munn, B.R., Zhou, N., Yang, P., Feng, J., Li, Y., Shine, J.M., Reimer, J., 2026. Inferring norepinephrine dynamics from partial observations reveals the temporal structure of elevations during arousal. bioRxiv 2026.03.29.715097. 10.64898/2026.03.29.715097

O’Toole, S.M., Oyibo, H.K., Keller, G.B., 2023. Molecularly targetable cell types in mouse visual cortex have distinguishable prediction error responses. Neuron 111, 2918–2928.e8. 10.1016/j.neuron.2023.08.015

Rajan, R., Dias, R.F., Malakasis, N., Baeta, M., Zhang, X., Gjorgjieva, J., Petreanu, L., 2026. Visual experience exerts an instructive role on cortical feedback inputs to the primary visual cortex. Current Biology 36, 1033–1044.e7. 10.1016/j.cub.2026.01.031

Rao, R.P., Ballard, D.H., 1999. Predictive coding in the visual cortex: a functional interpretation of some extra-classical receptive-field effects. Nat. Neurosci. 2, 79–87. 10.1038/4580

Reimer, J., McGinley, M.J., Liu, Y., Rodenkirch, C., Wang, Q., McCormick, D.A., Tolias, A.S., 2016. Pupil fluctuations track rapid changes in adrenergic and cholinergic activity in cortex. Nat Commun 7, 13289. 10.1038/ncomms13289

Roth, M.M., Dahmen, J.C., Muir, D.R., Imhof, F., Martini, F.J., Hofer, S.B., 2016. Thalamic nuclei convey diverse contextual information to layer 1 of visual cortex. Nat Neurosci 19, 299–307. 10.1038/nn.4197

Sales, A.C., Friston, K.J., Jones, M.W., Pickering, A.E., Moran, R.J., 2019. Locus Coeruleus tracking of prediction errors optimises cognitive flexibility: An Active Inference model. PLOS Computational Biology 15, e1006267. 10.1371/journal.pcbi.1006267

Saravanan, V., Berman, G.J., Sober, S.J., 2020. Application of the hierarchical bootstrap to multi-level data in neuroscience. Neuron Behav Data Anal Theory 3, https://nbdt.scholasticahq.com/article/13927-application-of-the-hierarchical-bootstrap-to-multi-level-data-in-neuroscience.

Schneider, D.M., Sundararajan, J., Mooney, R., 2018. A cortical filter that learns to suppress the acoustic consequences of movement. Nature 561, 391–395. 10.1038/s41586-018-0520-5

Solyga, M., Keller, G.B., 2025. Multimodal mismatch responses in mouse auditory cortex. eLife 13, RP95398. 10.7554/eLife.95398

Su, Z., Cohen, J.Y., 2022. Two types of locus coeruleus norepinephrine neurons drive reinforcement learning. 10.1101/2022.12.08.519670

Tsukano, H., Garcia, M.M., Dandu, P.R., Kato, H.K., 2026. Orbitofrontal cortex drives predictive filtering of sensory responses. Nat Neurosci 1–13. 10.1038/s41593-026-02217-z

Warren, R.A., Zhang, Q., Hoffman, J.R., Li, E.Y., Hong, Y.K., Bruno, R.M., Sawtell, N.B., 2021. A rapid whisker-based decision underlying skilled locomotion in mice. eLife 10, e63596. 10.7554/eLife.63596

Widmer, F.C., O’Toole, S.M., Keller, G.B., 2022. NMDA receptors in visual cortex are necessary for normal visuomotor integration and skill learning. eLife 11, e71476. 10.7554/eLife.71476

Wright, W.J., Hedrick, N.G., Komiyama, T., 2025. Distinct synaptic plasticity rules operate across dendritic compartments in vivo during learning. Science 388, 322–328. 10.1126/science.ads4706

Yogesh, B., Keller, G.B., 2025. Activity in serotonergic axons in visuomotor areas of cortex is modulated by the recent history of visuomotor coupling. Peer Community J 5, pcjournal.592. 10.24072/pcjournal.592

Yogesh, B., Keller, G.B., 2024. Cholinergic input to mouse visual cortex signals a movement state and acutely enhances layer 5 responsiveness. eLife 12. 10.7554/eLife.89986.4

Yu, A.J., Dayan, P., 2005. Uncertainty, neuromodulation, and attention. Neuron 46, 681–692. 10.1016/j.neuron.2005.04.026

Zmarz, P., Keller, G.B., 2016. Mismatch Receptive Fields in Mouse Visual Cortex. Neuron 92, 766–772. 10.1016/j.neuron.2016.09.057

